# Transcriptome wide association study of coronary artery disease identifies novel susceptibility genes

**DOI:** 10.1101/2021.07.21.453208

**Authors:** Ling Li, Zhifen Chen, Moritz von Scheidt, Andrea Steiner, Ulrich Güldener, Simon Koplev, Angela Ma, Ke Hao, Calvin Pan, Aldons J. Lusis, Shichao Pang, Thorsten Kessler, Raili Ermel, Katyayani Sukhavasi, Arno Ruusalepp, Julien Gagneur, Jeanette Erdmann, Jason C. Kovacic, Johan L.M. Björkegren, Heribert Schunkert

## Abstract

Transcriptome-wide association studies (TWAS) explore genetic variants affecting gene expression for association with a trait. Here we studied coronary artery disease (CAD) using this approach by first determining genotype-regulated expression levels in nine CAD relevant tissues by EpiXcan in two genetics-of-gene-expression panels, the Stockholm-Tartu Atherosclerosis Reverse Network Engineering Task (STARNET) and the Genotype-Tissue Expression (GTEx). Based on these data we next imputed gene expression in respective nine tissues from individual level genotype data on 37,997 CAD cases and 42,854 controls for a subsequent gene-trait association analysis. Transcriptome-wide significant association (P < 3.85e-6) was observed for 114 genes, which by genetic means were differentially expressed predominately in arterial, liver, and fat tissues. Of these, 96 resided within previously identified GWAS risk loci and 18 were novel (CAND1, EGFLAM, EZR, FAM114A1, FOCAD, GAS8, HOMER3, KPTN, MGP, NLRC4, RGS19, SDCCAG3, STX4, TSPAN11, TXNRD3, UFL1, WASF1, and WWP2). Gene set analyses showed that TWAS genes were strongly enriched in CAD-related pathways and risk traits. Associations with CAD or related traits were also observed for damaging mutations in 67 of these TWAS genes (11 novel) in whole-exome sequencing data of UK Biobank. Association studies in human genotype data of UK Biobank and expression-trait association statistics of atherosclerosis mouse models suggested that newly identified genes predominantly affect lipid metabolism, a classic risk factor for CAD. Finally, CRISPR/Cas9-based gene knockdown of RGS19 and KPTN in a human hepatocyte cell line resulted in reduced secretion of APOB100 and lipids in the cell culture medium. Taken together, our TWAS approach was able to i) prioritize genes at known GWAS risk loci and ii) identify novel genes which are associated with CAD.

## Introduction

Coronary artery disease (CAD), a leading cause of premature death worldwide, is influenced by interactions of lifestyle, environmental, and genetic risk factors^1^. Genome-wide association studies (GWAS) have identified over 200 risk loci for CAD^2, 3^. Most of them are located in non-coding regions which hampers their functional interpretation. Expression quantitative traits loci (eQTLs) to some extent explain the genomic effects of GWAS signals^4–6^. By leveraging effects of multiple *cis*-eQTL variants on gene expression, transcriptome-wide association studies (TWAS) search primarily for gene-trait associations. The approach builds on predictive models of gene expression derived from reference panels that correlate genotype patterns with transcript levels in tissues which are relevant for the phenotype. Predictive models are then used to associate tissue-specific gene expression based on genotypes with a given trait in individuals of GWAS cohorts^7^. Since TWAS signals reflect gene expression levels, the approach can be used to prioritize candidate genes across disease-relevant tissues. Thereby, TWAS may point to causal genes at risk loci identified by GWAS and thus provide further insights on biological mechanisms^8, 9^. Moreover, TWAS increase the sensitivity to identify susceptibility genes missed by traditional GWAS analyses. Here we performed a TWAS to identify novel susceptibility genes for CAD comprising more than 80,000 individuals with genotype data along with validation and exploratory analyses for the associated genes.

## Results

### Evaluation of the predictive models from STARNET and GTEx panels

The study design is shown in Fig. 1. We applied predictive models of nine tissues trained by the EpiXcan pipeline^9^ from two genetics-of-gene-expression panels: Stockholm-Tartu Atherosclerosis Reverse Network Engineering Task (STARNET) and Genotype-Tissue Expression (GTEx)^10, 11^. STARNET is a genetics-of-gene-expression study on approximately 600 CAD patients undergoing open-heart surgery, during which seven tissues were collected: atherosclerotic aortic wall (AOR), atherosclerotic-lesion-free internal mammary artery (MAM), liver (LIV), blood (BLD), subcutaneous fat (SF), visceral abdominal fat (VAF), and skeletal muscle (SKLM)^10^. GTEx is a comprehensive resource for genetics-of-gene-expression across 54 non-diseased tissue sites obtained post-mortem from nearly 1000 individuals^11^. In GTEx we studied six of the above tissues as well as the wall of coronary (COR) and tibial (TIB) arteries, whereas MAM was not available (Methods and Supplementary Tables 1-2). Together, we obtained predictive models from nine CAD-relevant tissues. Genes with cross-validated prediction R2 > 0.01 were kept. STARNET-based models allowed to impute 12,995 unique gene expression signatures in seven tissues, and GTEx 12,964 unique gene expression signatures in eight tissues (Supplementary Table 1).

**Fig. 1.**
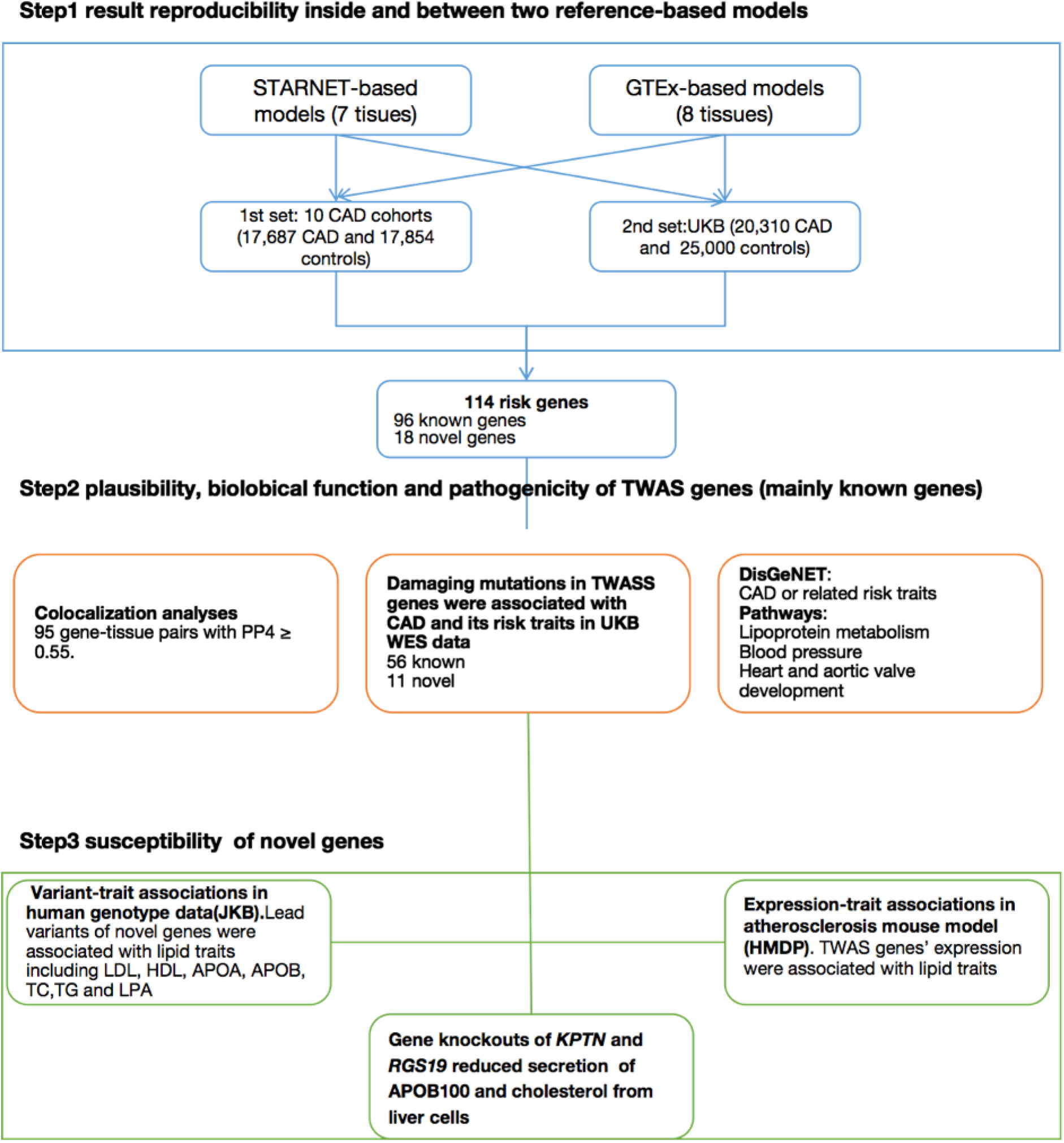
The study design.

We first tested the reproducibility of the STARNET- and GTEx-based predictive models by performing TWAS analyses in ten GWAS studies of CAD covering 17,687 CAD patients and 17,854 controls^12–21^, which provided individual level data and partially overlap with the CARDIoGRAMplusC4D meta-analysis, followed by replication analyses on genotyping data of UK Biobank (UKB)^22^, from which we extracted 20,310 CAD patients and 25,000 controls (Supplementary Table 3). As can be seen in Supplementary Results, there were prominent overlaps of transcriptome-wide significant genes having consistent association directions between test and validating sets within STARNET-(binomial test P = 0.00075) and GTEx-based models (binomial test P = 0.00079; Supplementary Fig. 1) respectively. Between the two independent reference panels, TWAS results of six overlapping tissues indicated consistent association directions (average Pearsońs coefficient ρ = 0.72; P < 1e-10; Supplementary Fig. 2), and prominent overlaps of significant gene-tissue pairs (Supplementary Results; Supplementary Fig. 3). Overall, these results suggest the reproducible of TWAS results of predictive models within and between two independent reference panels.

### Genes associated with CAD by TWAS

By combining TWAS results based on two genetics-of-gene-expression reference panels, we identified 114 genes representing 193 gene-tissue pairs with differential expression in CAD cases and controls (Fig. 2; Supplementary Fig. 4; Supplementary Table 4). Moreover, 95 of overall 114 gene-tissue association pairs were confirmed using another commonly used fine-mapping tool (COLOC)^23^ that calculates the posterior probabilities of shared casual variant in each locus between eQTL and GWAS statistics (Methods; Supplementary Table 5; Supplementary Fig. 5).

**Fig. 2.**
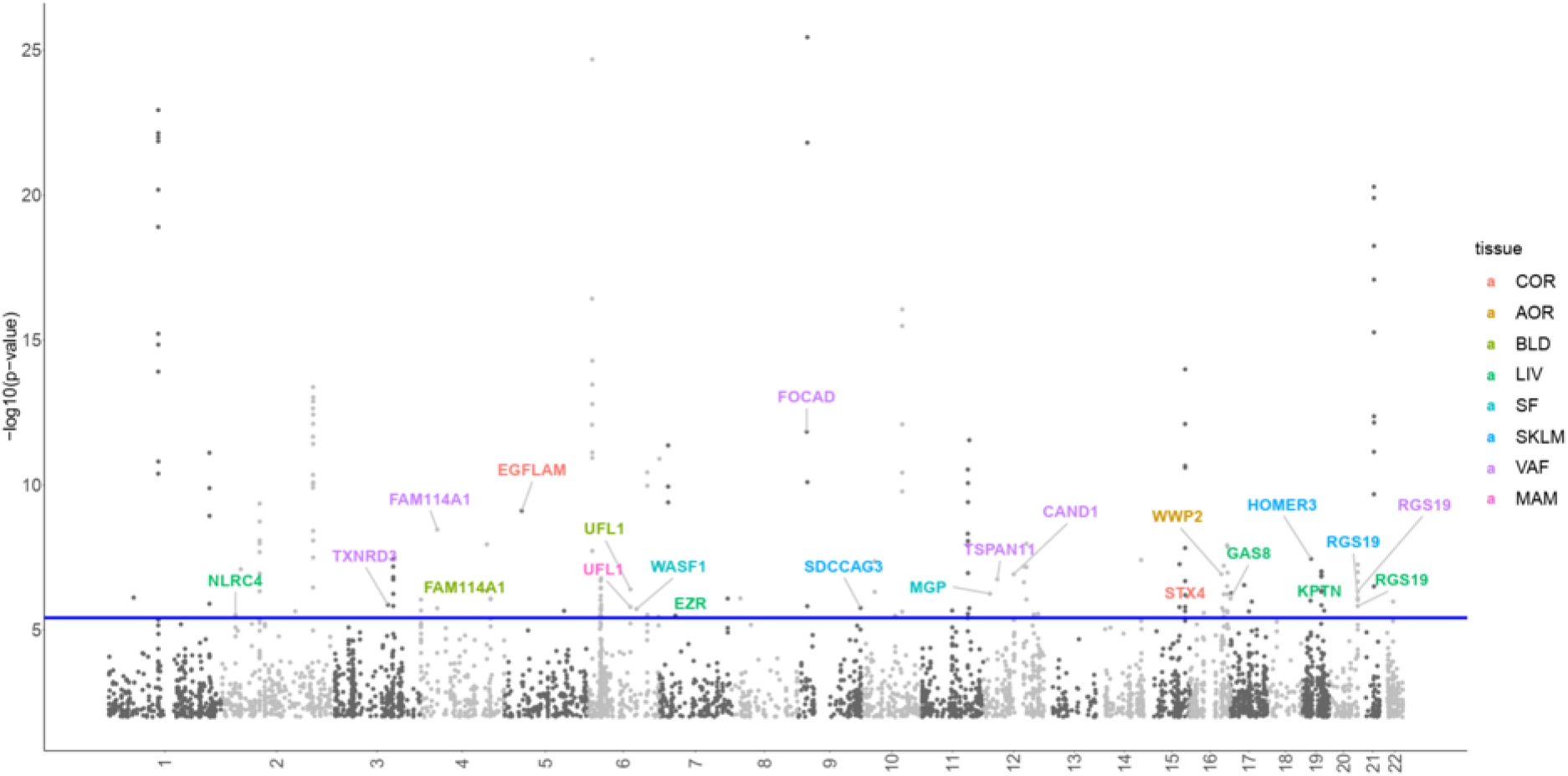
Manhattan plot of the transcriptome wide association study (TWAS). The results from STARNET- and GTEx-based TWASs were integrated by lowest P values. The blue line marks P =3.85-6. Each point corresponds to an association test between gene-tissue pair. 18 novel TWAS genes were highlighted. Supplementary Fig. 4 identifies all genes identified by their genetically-modulated association signals.

Forty-six genes displayed genetically-mediated differential expression in AOR, 28 in MAM, 25 in LIV, 23 in VAF, 22 in SKLM, 18 in SF, 16 in BLD, 10 in TIB, and 5 in COR (Fig. 3A), reflecting the importance of respective tissues in CAD pathophysiology. Most genes revealed significant associations in only a single tissue; 38 were significant in more than one, almost all having consistent directions of association between predicted expression levels and CAD across tissues (Fig. 3B).

**Fig. 3.**
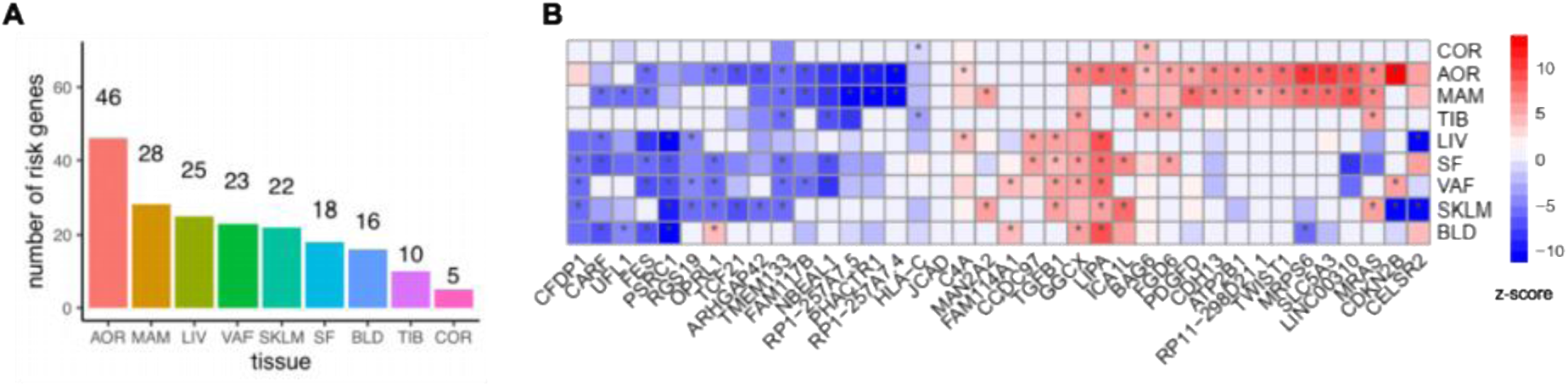
Tissue distribution of 114 CAD TWAS genes. (A) Number of significant genes across tissues. (B) Heatmap plot of 38 TWAS genes identified in more than one tissues. The color codes indicate direction of effects. Cells marked with * represent significant gene-tissue pairs (P < 3.85e-6).

Among the 114 genes, 102 were protein-coding and 12 were long non-coding RNAs (lncRNA) (Supplementary Table 4). STARNET data showed that most lncRNAs were positively co-expressed with a surrounding gene in affected tissues (Supplementary Fig. 8). *LINC00310* was the only exception, which displayed complex co-expression patterns with other genes (Supplementary Fig. 8).

Respective genes were found in 63 genomic regions, thus several regions represented multiple genes with significant associations. Six regions had multiple TWAS genes with shared GWAS and eQTL signals in respective tissues, like 1p13.3 and 2p33.2 (Supplementary Fig. 6-7; Supplementary Table 5). On the other hand, in 39 regions expression of only a single gene was found to be significantly associated, which makes these genes likely candidates for mediating causal effects, particularly, if these genes reside within GWAS risk loci for CAD (these genes are indicated in Supplementary Table 6).

Most TWAS genes (n=96) could be positionally annotated to the 1Mb region around one of the over 200 GWAS loci that are currently known to be genome-wide significantly associated with CAD^2, 3^. Therefore we marked these as known genes (Supplementary Table 6). On the other hand, 18 genes resided outside of these regions and were labeled as novel genes (Table 1). Most novel genes were tissue-specific, except *RGS19*, *FAM114A1* and *UFL1* which displayed evidence for differential expression in multiple tissues.

**Table 1.**
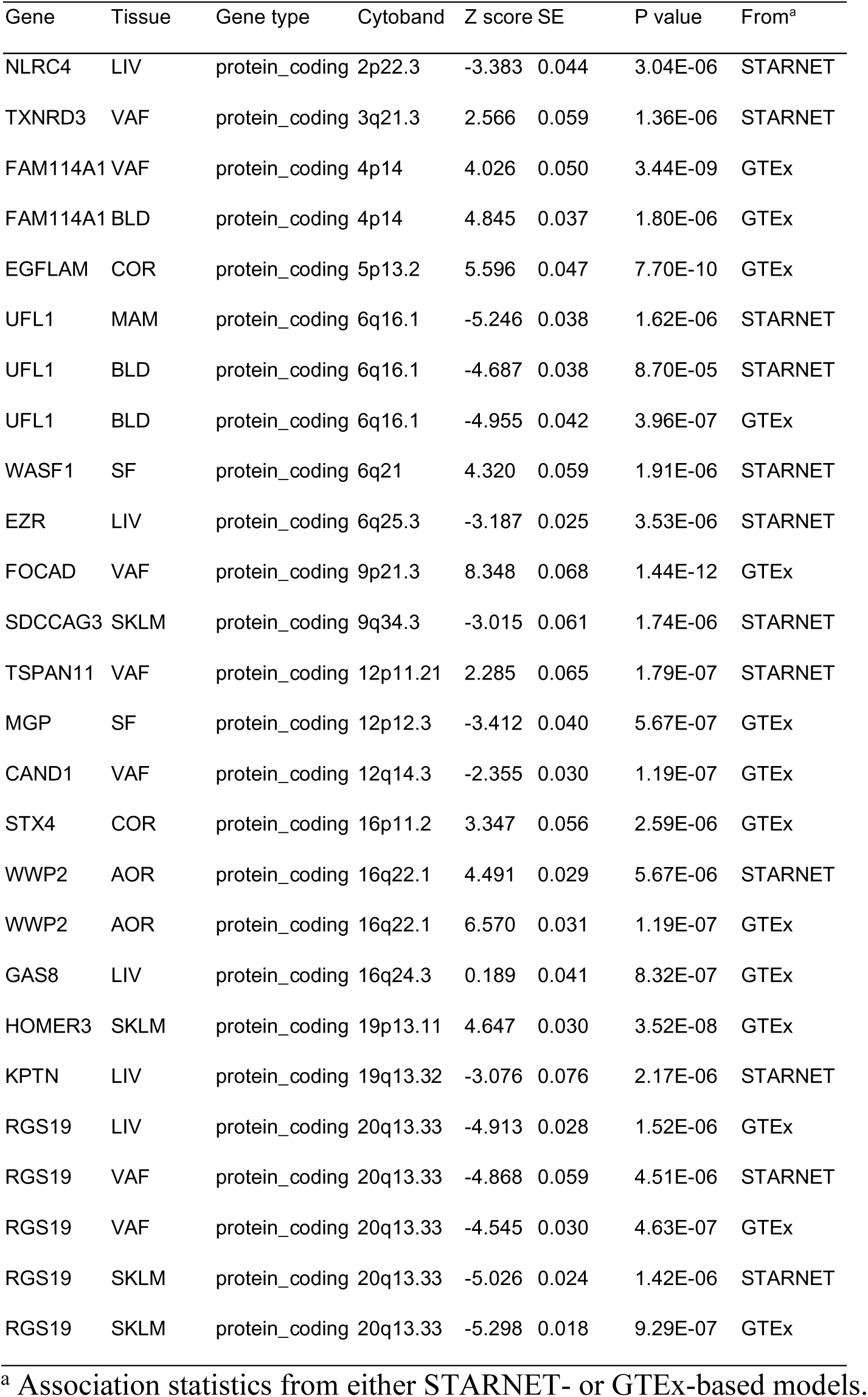
18 TWAS genes residing outside of published GWAS loci.

### Pathways and diseases enriched by TWAS genes

We carried out two types of gene set enrichment tests to further study the biological relevance of genes giving signals in this TWAS. First, we studied disease-gene sets from the DisGeNET platform which is one of the largest publicly available collections of genes and variants associated with human diseases^24^. The results showed that genes discovered by TWAS were primarily enriched for CAD, coronary atherosclerosis, and hypercholesterolemia (Supplementary Table 7), adding to the plausibility of our TWAS findings.

In line with these results, gene set enrichment analyses using GO^25^, KEGG^26^, Reactome^27^, and WikiPathways^28^ databases showed that the TWAS genes were highly enriched for pathways involved in cholesterol metabolism and regulation of lipoprotein levels. To a lesser extent, risk genes were enriched in regulation of blood pressure as well as development and morphogenesis of the heart and the aortic valve (Supplementary Table 8).

### Damaging mutations in TWAS genes

We next searched in whole-exome sequencing data of 200,643 participants from UKB for rare damaging variants in TWAS genes (minor allele frequency < 0.01, either loss of function mutations or mutations predicted to be adverse by one of five in-silico methods (Supplementary Files). We performed gene-based burden test on major CAD-related cardiometabolic risk traits. We found evidence for nominally significant association with either CAD or its risk traits for 67 TWAS genes (Fig. 4; Supplementary Tables 9-10). Mutations in five genes were directly associated with increased CAD risk: *LPL* (odds ratio [OR] = 1.168; 95% confidence interval [CI] 1.034-1.036; P = 0.016), *NOS3* (OR = 1.143; 95% CI 1.109-1.279; P = 0.02), *ADAMTS7* (OR = 1.062; 95% CI 1.011-1.115; P = 0.016), *MTAP* (or=1.507; 95%CI 1.061-2.086; P = 0.017), and *HLA-C* (OR = 1.112; 95%CI 1.002-1.239; P = 0.044); and two were associated reduced CAD risk: *TWIST1* (OR = 0.726; 95% CI 0.523-0.985; P = 0.038), *SARS* (OR = 0.831; 95% CI 0.706-0.974; P = 0.022). Damaging *LPL* mutations were evidently associated with lipid traits, including levels of LDL (low density lipoproteins) (beta = 0.043; P = 9.6e-4), HDL (high density lipoproteins) (beta = - 0.106; P = 4.54e-68), APOA (Apolipoprotein A) (beta = -0.062; P = 6.25e-47), APOB (Apolipoprotein B) (beta = 0.025; P = 1.38e-12), and TG (Triglycerides) (beta = 0.241; P = 1.47e-68).

**Fig. 4.**
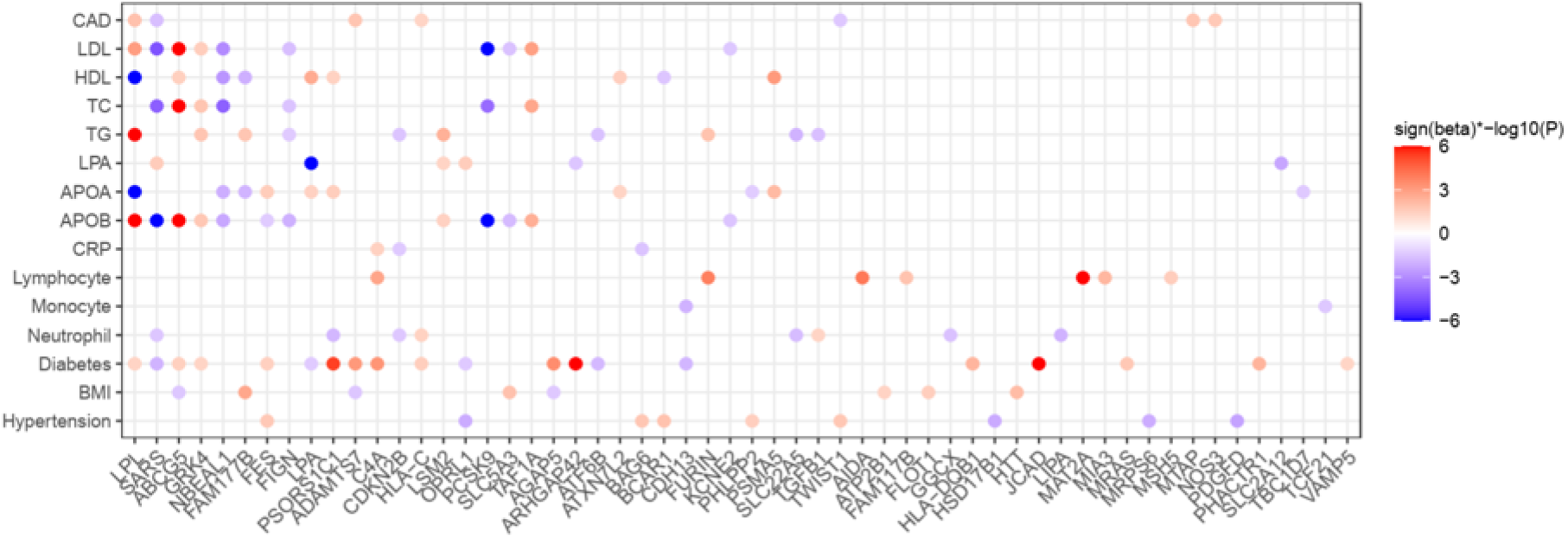
Effects of damaging mutations of TWAS genes on CAD and its risk traits. Sign(beta)*-log10(p) displayed for associations that reached a P <0.05. When the Sign(beta)*-log10(P) > 6, they were trimmed to 6

Damaging mutations in 11 novel TWAS genes were associated with CAD risk factors (Table 2). Some of these gene-trait associations have been reported before. Damaging mutations in *MGP*, which regulates vascular calcification, adipogenesis and is serum marker of visceral adiposity^29–31^, were associated with increased levels of LDL, TC (total cholesterol) and APOB. *NLRC4* was reportedly associated with atherosclerosis by regulating inflammation reaction^32, 33^, and its damaging mutations were associated with levels of CRP (C-reactive protein – a marker of inflammation).

**Table 2.** Associations of damaging mutations in novel genes with risk traits of CAD.

| Binary trait | Gene | Case |  | Control |  | OR[95%CI] | P value |
| --- | --- | --- | --- | --- | --- | --- | --- |
|  |  | Non-carrier | Carrier | Non-carrier | Carrier |  |  |
| Diabetes | FAM114A1 | 10668 | 116 | 187555 | 1457 | 1.4[1.15-1.69] | 9.19E-04 |
| Diabetes | UFL1 | 10634 | 150 | 187023 | 1989 | 1.33[1.11-1.57] | 1.47E-03 |
| Hypertension | FOCAD | 73542 | 4605 | 102379 | 6129 | 1.05[1.01-1.09] | 2.60E-02 |
| Hypertension | EGFLAM | 73754 | 4393 | 102147 | 6361 | 0.96[0.92-1] | 2.82E-02 |
| Hypertension | EZR | 77495 | 652 | 107491 | 1017 | 0.89[0.8-0.98] | 2.05E-02 |
| Quantitative trait | Gene | Carrier |  | Non-carrier |  | Beta[95%CI] | P value |
|  |  | No. carrier | Median(range) | No. non-carrier | Median (range) |  |  |
| APOB (g/L) | HOMER3 | 2633 | 1(0.41-1.91) | 187891 | 1.02(0.4-2) | -0.02[-0.03--0.01] | 4.02E-03 |
| APOB (g/L) | MGP | 158 | 1.05(0.51-1.96) | 190366 | 1.02(0.4-2) | 0.08[0.04-0.13] | 2.60E-04 |
| TC (mmol/L) | HOMER3 | 2651 | 5.57(2.33-10.06) | 188814 | 5.66(1.64-15.46) | -0.08[-0.14--0.03] | 2.95E-03 |
| TC (mmol/L) | MGP | 158 | 5.76(3.19-10.29) | 191307 | 5.66(1.64-15.46) | 0.34[0.13-0.56] | 1.66E-03 |
| LDL (mmol/L) | HOMER3 | 2649 | 3.45(1.05-6.97) | 188511 | 3.52(0.28-9.8) | -0.06[-0.11--0.02] | 2.34E-03 |
| LDL (mmol/L) | MGP | 158 | 3.59(1.81-7.05) | 191002 | 3.52(0.28-9.8) | 0.29[0.13-0.45] | 4.82E-04 |
| LPA (nmol/L) | TXNRD3 | 3162 | 21.94(3.8-188.89) | 150645 | 20.98(3.8-189) | 2.5[0.29-4.71] | 2.63E-02 |
| BMI (kg/m2) | KPTN | 2084 | 26.87(14.94-56.05) | 197753 | 26.7(12.12-68.95) | -0.3[-0.57--0.04] | 2.65E-02 |
| BMI (kg/m2) | WASF1 | 806 | 26.92(17.71-53.02) | 199031 | 26.7(12.12-68.95) | 0.47[0.04-0.91] | 3.38E-02 |
| CRP (mg/L) | NLRC4 | 2470 | 1.25(0.11-52.86) | 188577 | 1.31(0.08-79.49) | -0.22[-0.44--0.01] | 4.30E-02 |
| CRP (mg/L) | UFL1 | 2057 | 1.3(0.1-43.74) | 188990 | 1.31(0.08-79.49) | -0.37[-0.6--0.13] | 2.36E-03 |
| Neutrophil (10<sup>9</sup> cells/L) | MGP | 164 | 3.51(0.61-8.21) | 194782 | 4.07(0-25.95) | -0.33[-0.59--0.07] | 1.40E-02 |

### Novel genes associate with risk factors in human and mouse data

We next associated common variants in the regions of ±1Mb around the 18 novel TWAS genes to study their associations with a series of lipid traits including LDL, HDL, APOA, APOB, LPA, TC, and TG in UKB (Supplementary Files). Bonferroni-corrected significance P<4.0e-4 (0.05/18 novel genes * 7 lipid traits) was observed for numerous respective lead variants, of which *RGS19*, *SDCCAG3*, *HOMER3*, and *WWP2* reached genome-wide significant association (P<5e-8) with multiple lipid traits (Fig. 5A; Supplementary Table 11).

**Fig. 5.**
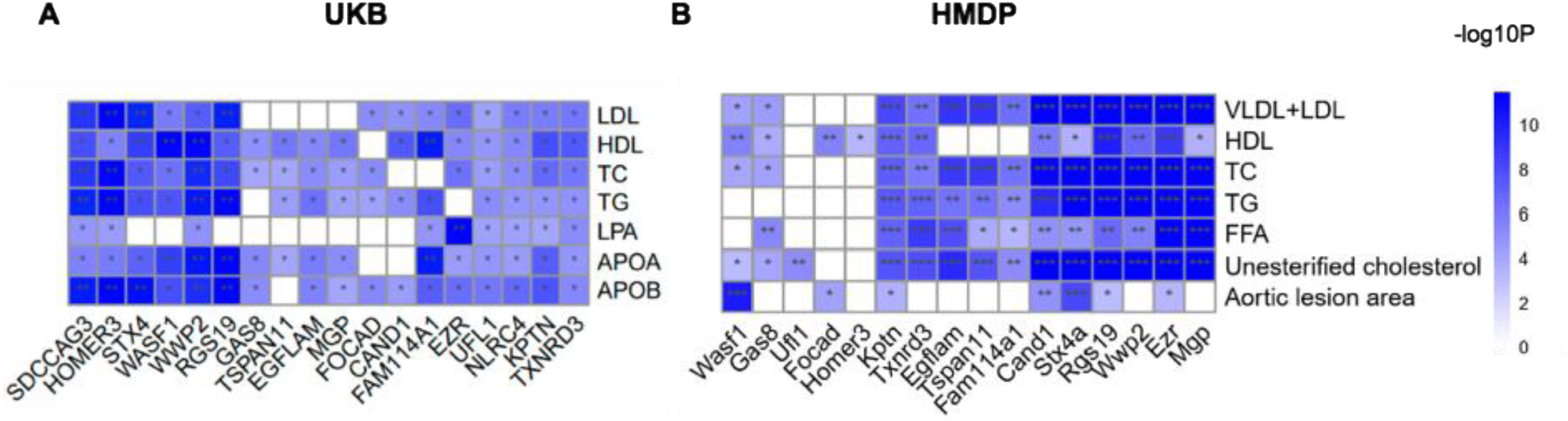
Novel risk genes were associated with lipid traits. (A) Data from UKB indicate that lead variants inside the boundary of risk genes were associated with lipid traits with Bonferroni-corrected significance levels (*, P < 4.0e-4), or by genome-wide significance (**, P < 5e-8). (B) Expression levels of novel genes were likewise associated with lipid traits and aortic lesion area in an atherosclerosis mouse model from the Hybrid Mouse Diversity Panel (HMDP). *, P <0.05; **, P<0.01; ***, P < 0.001.

Next, we extracted expression-trait association statistics of TWAS genes from the Hybrid Mouse Diversity Panel (HMDP)^34^. Based on the expression data from mouse aorta and liver tissues, 48 TWAS genes were significantly associated with aortic lesion area and 14 further cardiovascular traits (nominal significance P < 0.05; Supplementary Table 12). Expression levels of seven novel genes, i.e. *Rgs19*, *Kptn*, *Ezr*, *Stx4a*, *Cand1*, *Focad* and *Wasf1,* were associated with aortic lesion area (Fig. 5B), a commonly used measure for atherosclerotic plaque formation in mice. Additionally, we found the novel genes were associated with at least one lipid trait in the mouse model (Fig. 5B).

### Knockdown of *RGS19* and *KPTN* reduced lipid secretion by human liver cells

Both human genotype-trait association statistics in UKB and mouse expression-trait association statistics in the HMDP indicated that several novel genes identified by TWAS influence lipid metabolism. To validate these findings, we chose two of the novel genes, i.e. *KPTN* and *RGS19*, which have not been studied in much detail so far and have particularly not at all been investigated in the context of atherosclerosis or CAD. Hepatocytes are critically involved in lipid metabolism. In line, in a screening of different atherosclerosis-relevant cell lines (e.g., hepatocytes, smooth muscle, endothelium, fibroblast, and adipocytes), *KPTN* had the highest expression level in the huh7 hepatocyte cell line (Supplementary Fig. 9A, B). To study the influence of *KPTN* and *RGS19* on lipid metabolism, we next generated gene knockout (KO) huh7 cell lines for by a dual CRISPR strategy (Methods; Supplementary Table 13), which substantially reduced expression of the respective genes (Supplementary Fig. 9C, D). We measured secretion levels of TG, cholesterol and APOB in gene-targeted versus control cells. Notably, under normal circumstances, human hepatocytes synthesize cholesterol, assemble TG and APOB100, and secrete these particles in form of very low-density lipoprotein (VLDL)^35^. Compared to control huh7 cells, we found reduced APOB and cholesterol levels in culture medium of *KPTN*-KO cells (Fig. 6C, D). In culture medium of *RGS19*-KO cells we also detected reduced levels of APOB100, cholesterol, and TG (Fig. 6B, C, D, E), in line with strong associations of this gene with an array of lipid traits in both human genotyping and mouse expression data sets (Figure 5).

**Fig. 6.**
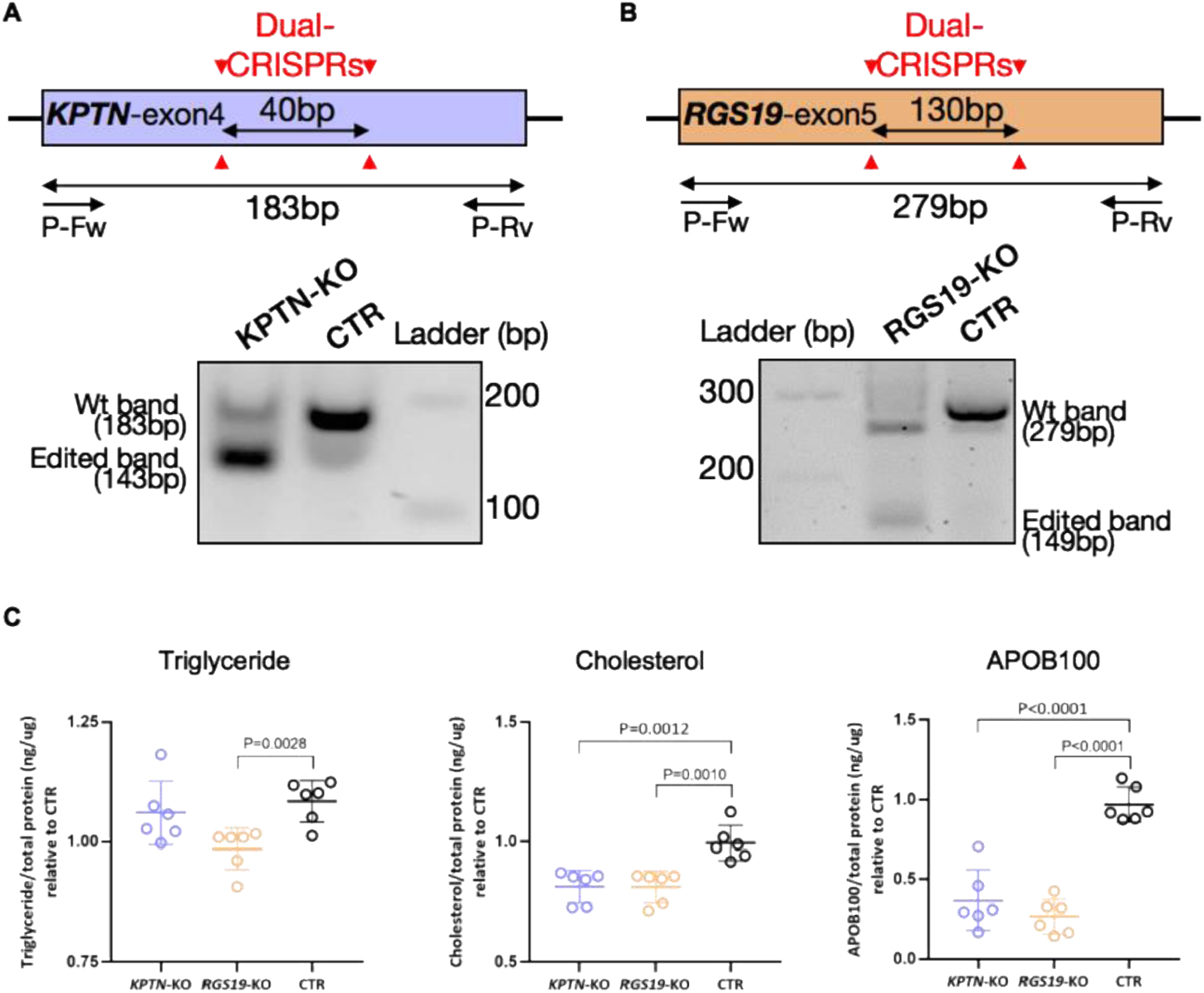
Targeting of *KPTN* and *RGS19* reduced Lipids and APOB secretion of human liver cells. (A) Two sgRNAs were used to target the exon4 of *KPTN* (shared exon among isoforms) in a Cas9-expressing huh7 liver cell line. The dual CRISPR strategy created a 40bp frame shift deletion in the gene and pround reduction of *KPTN* at both mRNA and protein levels (Supplementary Figure 9C, 9D). The primers (P-Fw and P-Rv) used for analyzing the CRISPR editing as indicated. (B) The same strategy was used for *RGS19* targeting, which resulted in a 130bp frame shift deletion in the gene, and reduction of mRNA and protein (Supplementary Figure 9C, 9D). (C) Reduced triglyceride and cholesterol levels in knockout (KO) cell lines were detected by colorimetric method and APOB100 secretion was measured by human APOB100 Elisa (n=6). Triglyceride, cholesterol and APOB100 levels were normalized to total protein and compared between the KO and control (CTR) cell lines.

## Discussion

In a stepwise approach, we first generated and filtered models predicting genetically modulated gene expression in nine tissues that contribute to CAD risk. Next, we applied these models to individual-level genotype data on more than 80,000 CAD cases and controls. We identified 114 genes with differential expression by genetic means in CAD patients. Many signals were highly plausible as they resided within loci displaying genome-wide significant association with CAD by traditional GWAS. Moreover, the genes identified by this TWAS were markedly enriched in established pathways for the disease, and 67 revealed in whole-exome sequence data of UKB that damaging mutations have significant impact on CAD risk or its underlying traits. Importantly, we also identified 18 genes without prior evidence for their involvement in CAD by GWAS, many of which were found to be associated with lipid metabolism in human and mouse data.

Only a minority of genes residing within published CAD GWAS loci have been validated experimentally for their underlying causal role in atherosclerosis. Our data corroborate a recent exploration of known GWAS loci for genotype-related expression levels (Hao et al., personal communication, manuscript attached) and provide a substantial step towards prioritization of genes at respective GWAS loci^2, 3^. In this respect, 46 genes identified by this TWAS are known for effects in pathophysiological pathways related to CAD, including lipid metabolism, inflammation, angiogenesis, transcriptional regulation, cell proliferation, NO signaling, and high blood pressure, to name a few (Supplementary Table 6), giving credibility to the association findings. On the other hand, a limitation of the TWAS approach is that at 20 loci two or more genes show signals such that other methods will be needed to pinpoint the precise genetic mechanisms leading to CAD. Indeed, in another study we recently applied summary-based Mendelian Randomization, MetaXcan, to integrate tissue and cell-specific data from STARNET and GTEx with CAD GWAS datasets, and obtained at 14 of these 20 loci indicative data allowing prioritization of a gene (Hao et al., personal communication, manuscript attached).

Most novel TWAS genes revealed association with lipid traits in both genotype data of human and expression-trait statistics of our atherosclerosis mouse model. For example, expression profiles of *KPTN* and *RGS19*, both novel genes displaying significant TWAS results for CAD in human liver tissue, also showed significant association with various lipid traits as well as aortic lesion area in our atherosclerosis mouse model. Moreover, both gene loci harbor SNPs which are genome-wide significantly associated with LDL-C, HDL-C, TC, and TG in human genotype data. Finally, the Common Metabolic Disease Knowledge Portal revealed that damaging rare variants of *KPTN* are associated with reduced levels of LDL (beta = -11.9; P = 0.00042) and TC (beta = -11.9; P = 0.0014) ^36^, which is directionally plausible given the TWAS results. Based on these observations, we functionally validated the roles of these two novel genes by studying lipid levels in human liver cells, i.e. the tissue that displayed evidence for differential expression by TWAS. Indeed, we observed that knockout of these genes lowered secretion of APOB and cholesterol into culture medium. *KPTN*, kaptin (actin binding protein), a member of the *KPTN*, *ITFG2*, *C12orf66* and *SZT2* (KICSTOR) protein complex, is a lysosome-associated negative regulator of the mechanistic target of rapamycin complex 1 (mTORC1) signaling^37^. It is required in amino acid or glucose deprivation to inhibit cell growth by suppressing mTORC1 signaling in liver, muscle, and neurons. mTORC1 has multifaceted roles in regulating lipid metabolism, including the promotion of lipid synthesis, and storage and inhibition of lipid release and consumption, suggesting that the validated role of *KPTN* in hepatic lipid secretion might be partially mediated by the mTORC1 pathway.*RGS19* belongs to the *RGS* (regulators of G-protein signaling) family, who are regulators for G protein-coupled receptors (GPCRs)^38^. *RGS19* inhibits GPCR signal transduction by increasing the GTPase activity of G protein alpha subunits, thereby transforming them into an inactive GDP-bound form^39, 40^. The targeting GPCR of RGS19 has not been observed before, and how RGS19 regulates lipid metabolism remains unclear.

Interestingly, our TWAS uncovered eight novel gene-CAD associations in fat tissue, including *MGP* and *WASF1* in SF, and *CAND1*, *FAM114A1*, *FOCAD*,*RGS19*,*TSPAN11* and *TXNRD3* in VAF, representing half of the novel genes. Damaging mutations in five genes were associated with many cardiometabolic risk factors for CAD, including those in *WASF1* with BMI, *MGP* with LDL,TC and APOB, *TXNRD3* with LPA, *FAM114A1* with diabetes, *FOCAD* with hypertension, i.e. conditions shown by Mendelian randomization to be causal for CAD^41^. Given the many CAD patients that are overweight or obese, it will be of great interest to identify how these genes modify cardiometabolic traits leading to cardiovascular disorders. In this respect our TWAS could provide a list of candidate genes and related targetable cardiometabolic traits. In addition, it is of surprise to unveil 22 genes linking SKLM to CAD risk, and eight were unique to this tissue, including *HOMER3*, *SDCCAG3*, *MTAP*, *NME9*, *PSMA4*, *SLC2A12*, *UNC119B* and *VAMP5*, the first two being novel. *SDCCAG3* or *ENTR1* encodes endosome associated trafficking regulator 1 and involves in recycling of *GLUT1* (glucose transporter type 1), supplying the major energy source for muscle contraction. SKLM-based metabolism may have a protective role in CAD as suggested by the many cardioprotective effects of sports^42, 43^. Gene targets enhancing SKLM function in this respect might be effective in CAD prevention, a field relatively unexplored thus far. Here, for the first time, quantitative traits regulated genes in SKLM were associated with CAD by TWAS, providing novel evidence for genes that could modulate CAD risk by their functions in SKLM.

There are certain limitations in our study. Since TWAS are strongly dependent on the reference panel linking genetic signatures with gene expression, it had to be expected that STARNET- and GTEx-based predictive models display differences in gene-CAD associations. STARNET-based TWAS identified 86 genes, whereas GTEx-based TWAS identified 68 genes. Yet, 34 genes were shared between the two analyses, and effect sizes for the shared genes were highly concordant (ρ = 0.97). An average of 62% overlapping genes was observed in the matched tissues of two reference-based models, and the resulting size of expression-CAD associations was linearly consistent with an average ρ = 0.72. The relatively small differences may be due to different sample sizes used for training predictive models^9^, different disease states (subjects with and without CAD), intravital or *post mortem* sample collection, leading to differences in gene expression in our reference panels^10, 11^. Given a fair consistency between the two data sources, we combined results derived from both panels to increase the power for capturing risk genes. Second, although TWAS facilitates candidate risk gene prioritization, co-regulation or co-expression *in cis* at a given locus limits the precise determination of the culprit gene^8^. Indeed, at 12 loci we observed signals for three or more TWAS genes. For instance, in LIV tissue TWAS identified five genes at 1p13.3, *ATXN7L2*, *CELSR2*, *PSMA5*, *PSRC1*, *SARS* and *SORT1* which were co-regulated by same risk variant set, confusing the causal gene prioritization. While *CELSR2*, *PSRC1* and *SORT1* were previously shown to act on lipid metabolism^44^, we found that damaging mutations in *ATXN7L2* and *SARS* were also associated with CAD or its risk traits, the former with serum levels of HDL and APOA, and the later with CAD and diabetes. In addition, all lncRNA genes identified by our study displayed co-expression with their neighboring coding genes, which makes it difficult to determine their casual effects. Nevertheless, in combining TWAS data with other genetic analyses, e.g., looking at effects of damaging mutations, genetic association with other phenotypes and expression-traits association statistics, we aimed to improve risk gene prioritization, and to provide deeper insights of possible disease-causing mechanisms. For instance, *LPL* is well-known for its protective role against CAD by lowering lipids^45, 46^, and our analyses showed that damaging *LPL* mutations were associated with increased risk of CAD and higher lipid levels. Finally, as with all statistical methods, there are certain limitations and assumptions associated with TWAS. Further evolution and improvement of these methods, as well as functional validation experiments, will assuredly improve the accuracy of these studies.

In summary, our TWAS study based on two genotype-expression reference panels identified 114 gene-CAD associations, of which 18 were novel. The extended analyses with multiple datasets supported the reliability of the CAD TWAS signals in prioritizing candidate risk genes and identifying novel associations in a tissue-specific manner. Functional validation of two novel genes, *RGS19* and *KPTN,* lend support to our TWAS findings. Our study created a set of gene-centered and tissue-annotated associations for CAD, providing insightful guidance for further biological investigation and therapeutic development.

## Methods

### Predictive models of nine tissues based on two reference panels

We adopted the existing predictive models trained using EpiXcan pipeline by Zhang et al.^1^, including models of atherosclerotic aortic wall (AOR), atherosclerotic-lesion-free internal mammary artery (MAM), liver (LIV), blood (BLD), subcutaneous fat (SF), visceral abdominal fat (VAF) and skeletal muscle (SKLM) based on the genetics-of-gene-expression panel STARNET (The Stockholm-Tartu Atherosclerosis Reverse Network Engineering Task)^2^, and of AOR, LIV, BLD, SF, VAF and SKLM based on GTEx (Genotype-Tissue Expression)^3^.

Arterial wall coronary (COR) and tibial artery (TIB), datasets were only available in the GTEx panel. So, we established predictive models for these two tissues using EpiXcan pipeline as has been done for other models before^1^. In brief, we firstly filtered the genotype and expression data of COR and TIB from GTEx v7. Variants with call rate < 0.95, minor allele frequency (MAF) < 0.01, and Hardy Weinberg equilibrium (HWE) < 1e-6 were removed. For expression, we used quality-controlled data and performed sample-level quantile normalization, and gene-level inverse quantile normalization using preprocess codes of PredicDB pipeline. Samples were restrained to the European ethnicity. We then calculated SNP priors by using hierarchical Bayesian model (qtlBHM)^4^ that jointly analyzed epigenome annotations of aorta derived from Roadmap Epigenomics Mapping Consortium (REMC)^5^, and eQTL statistics. The SNP priors (Supplementary Table 2), genotype data and expression data were jointly applied to 10-fold cross-validated weighted elastic-net to train predicting models by deploying EpiXcan pipeline^1^.

Both STARNET- and GTEx-based models were filtered by cross-validated prediction R^2^ > 0.01. The summary statistics of sample sizes used for training models and the transcript numbers of genes covered by each predicting models are shown in Supplementary Table 1.

### Genotype cohorts

For the discovery cohort, individual level genotyping data were collected from ten genome-wide associations studies (GWAS) of coronary artery disease (CAD), a subset of CARDIoGRAMplusC4D, including the German Myocardial Infarction Family Studies (GerMIFS) I-VII^6–12^, Wellcome Trust Case Control Consortium (WTCCC)^13^, LURIC study^14^ and Myocardial Infarction Genetics Consortium (MIGen)^15^. We used a part of individual-level data from UK Biobank (UKB) as the replication cohort^16^, by extracting 20,310 CAD cases according to hospital episodes or death registries as reported, and randomly selected 25,000 non-CAD UKB participants as controls. The detailed information about selection criteria of case and control were described at elsewhere^12^. In total, genotyping data of 37,997 cases and 42,854 controls were included in our transcriptome-wide association studies (TWAS) of CAD (Supplementary Table 3). The preprocessing steps of genotyping data are as previously^12^.

### Transcriptome wide association analysis

We applied predictive models to the eleven genotype cohorts to impute individual-level expression profiles of nine tissues, and performed transcriptome-wide association analysis between imputed expression and CAD. To test the reproducibility of TWAS results, we performed two types of validating tests: within and between two reference-based models. Firstly, we used ten GWAS cohorts as testing set and UKB as the validating set to test reproducibility within STARNET- and GTEx-based models respectively. Secondly, we compared the consistency of results between STARNET- and GTEx-based models of the six overlapping tissues using all genotype data.

### Co-expression network for lncRNA

We used RNA-seq data of STARNET^2^ to calculate expression correlations between long non-coding RNA (lncRNA) genes and protein coding genes in seven tissues. Co-expression pairs with absolute Pearson correlation coefficient larger than 0.4 were considered to be significant. The co-expression network was displayed by cytoscape^17^.

### Colocalization of the eQTL and GWAS signals

Colocalization analysis was performed using COLOC, a Bayesian statistical methodology that takes GWAS and eQTL data as inputs, and tests the posterior probabilities (PP4) of shared casual variant for each locus^18^. The summary statistics of GWAS meta-analysis were obtained from CARDIoGRAMplusC4D Consortium^11^, and the eQTL data of nine tissues from STARNET^2^ and GTEx^3^ respectively.

### Annotation of novel risk genes

Over 200 CAD loci were identified by GWAS^19, 20^. We used MAGMA^21^ to annotate the 114 TWAS genes and observed that 96 genes resided within ±1Mb around known CAD loci whereas 18 genes (novel loci) where located outside known GWAS risk loci, i.e. they were novel genes (Supplementary Table 6).

### Gene set enrichment analyses

Pathway enrichment analysis was carried out using ClueGO (v2.5.2)^22^, a plugin of cytoscape^17^, based on collated gene sets from public databases including GO^23^, KEGG^24^, Reactome^25^, and WikiPathways^26^. Gene sets with false discovery rate (FDR) by right-sided hypergeometric test less than 0.05 were considered to be significant.

Furthermore, we also studied the diseases or traits associated with risk genes by performing disease enrichment analysis based on DisGeNET^27^, the largest publicly available datasets of genes and variants association of human diseases. FDR < 0.05 was used for thresholding.

### Rare damaging variants association analysis

To investigate association of damaging variants in TWAS genes with CAD, we used whole exome sequencing (WES) data of 200,632 participants from UKB^28^. The WES data was processed following the Functional Equivalence (FE) protocol. We performed quality control on the WES data by filtering variants with calling rate < 0.9, variants with HWE < 1e-6. For the relevant traits, besides CAD, we considered several risk factors of the disease, including body mass index (BMI), diabetes, hypertension, levels of low density lipoproteins (LDL), high density lipoproteins (HDL), apolipoprotein A (APOA), apolipoprotein B (APOB), Lipoprotein(a) (LPA), total cholesterol (TC) and triglycerides (TG)), as well as inflammation related factors (C-reactive protein (CRP), lymphocyte count (Lymphocyte), monocyte count (Monocyte) and neutrophil count (Neutrophil).

We defined damaging mutations as i) rare mutations with MAF < 0.01; ii) annotated into following one of the 3 classes: loss-of-function (LoF) (stop-gained, splice site disrupting, or frameshift variants), variants annotated as the pathogenic in ClinVar^29^, or missense variants predicted to be damaging by one of five computer prediction algorithms (LRT score, MutationTaster, PolyPhen-2 HumDiv, PolyPhen-2 HumVar, and SIFT). The Ensembl Variant Effect Predictor (VEP)^30^ and its plugin loftee^31^, and annotation databases dbNSFP 4.1a^32^ and ClinVar (GRCh38)^29^ were used for annotating damaging mutations.

For each analysis, samples were classified into carriers or noncarriers of the gene’s damaging mutations. For binary traits, we used Fisher’s exact test to check if there was incidences difference of mutation carrying between case and controls. For the quantitative traits, we used linear regression model with adjustments of sex, first five principal components, and lipid medication status to investigate the associations between mutation carrying status and traits. We used nominal significance threshold (P < 0.05), given that coding variants are rather rare, and the case-control sample sizes were not balanced which might increase false negative rate. We used nominal significance threshold P < 0.05, because, at one hand, the case-control size was not balanced which might increase false negative rate, at the other hand, it’s an exploratory trial to investigate the potential biological relevance of TWAS genes.

### Association of variants resided in novel genes with lipid traits

For 18 novel risk genes, we performed association analysis for variants located in novel gene loci (±1Mbase) with lipid traits using genotyping data of UKB. The lipid traits include levels of LDL, HDL, APOA, APOB, LPA, TC and TG. The variants were filtered by MAF > 0.01, and imputation info score > 0.4. The association test was performed using PLINK2^33^ with adjustment of sex, first five principal components, and lipid medication status. The lead variants residing in gene loci with P value less than 4.0e-4 (0.05/18 risk genes * 7 lipid traits) were considered to be significant (Supplementary Table 11).

### The Hybrid Mouse Diversity Panel (HMDP)

The Hybrid Mouse Diversity Panel (HMDP) is a set of 105 well-characterized inbred mouse strains on a 50% C57BL/6J genetic background^34^. To specifically study atherosclerosis in the HMDP, transgene implementation of human APOE-Leiden and cholesteryl ester transfer protein was performed, promoting distinct atherosclerotic lesion formation^35^. A Western diet containing 1% cholesterol was fed for 16 weeks. Subsequently, gene expression was quantified in aorta and liver of these mice and lesion size was assessed in the proximal aorta using oil red O staining. Other 14 related traits were measured too, including liver fibrosed area, body weight, total cholesterol, VLDL (very low-density lipoprotein) + LDL, HDL,TGs, unesterified cholesterol, free fatty acid, IL-1b, IL-6, TNFa, MCP-1, and M-CSF. From HMDP, we extracted significant association pairs between TWAS genes and 15 risk traits by applying significance P < 0.05.

### Experimental validation of *KPTN* and *RGS19* in human cells

To knock down *KPTN* and *RGS19*, two sgRNAs targeting shared exons of all transcription isoforms were delivered by lentivirus into a Cas9-expression huh7, a human hepatoma cell line. Exon 4 of *KPTN* and exon 5 of *RGS19* were targeted by a dual CRISPR strategy to create a 40bp and 130bp frame shift deletion, respectively. SgRNAs were carried by Lenti-Guide-Puro vector (addgene, #52963) and infected cells were treated with 10ug/ml puromycin treatment for 3 days to eliminate the negative cell. Positive targeted cells were expanded in culture and passaged for assays. Cells for measurement of secretive triglycerides, cholesterol and APOB100 were cultured for 16 hours in serum-free medium. Medium triglycerides and cholesterol were enriched for five times by vacuum centrifuge and measured with colorimetric kits, triglyceride (cobas) and CHOL2 (cobas), respectively. The amount of medium APOB100 was measured with an ELISA kit (MABTECH).

## Author Contributions

H.S., L.L., Z.C., designed the study and wrote the manuscript. L.L. ran analyses. Z.C. and A.S. performed experiments. M.V.S, U.G., S.C.P., S.K., C.P. A.J.L., T.K., A.R.,J.A., J.G., K.H., J.C.K. and J.M.B. provided research data, technical support and gave conceptual advice.

## Tools and Data

EpiXcan pipeline: https://bitbucket.org/roussoslab/epixcan/src/master/, and predictive models based on STARNET and GTEx databases: http://predictdb.org/

PrediXcan pipeline: https://github.com/hakyim/PrediXcan.

qtlBHM: https://github.com/rajanil/qtlBHM

STARNET database: https://www.ncbi.nlm.nih.gov/projects/gap/cgi-bin/study.cgi?study_id=phs001203.v1.p1. Project ID: 13585.

GTEx database: https://www.ncbi.nlm.nih.gov/projects/gap/cgi-bin/study.cgi?study_id=phs000424.v8.p2. Project ID: 20848.

UK Biobank: https://www.ukbiobank.ac.uk/. Project ID: 25214

MAGMA: https://ctg.cncr.nl/software/magma

R package for colocalization analysis, coloc: https://cran.r-project.org/web/packages/coloc/vignettes/vignette.html

DisGeNET: https://www.disgenet.org/

CARDIoGRAMplusC4D Consortium: http://www.cardiogramplusc4d.org/

## Extended data

## Supplementary Results

We tested the reproducibility of the STARNET- and GTEx-based predictive models by performing TWAS analyses in ten GWAS studies of CAD covering 17,687 CAD patients and 17,854 controls^12–21^, which provided individual level data and partially overlap with the CARDIoGRAMplusC4D meta-analysis, followed by replication analyses on genotyping data of UK Biobank (UKB)^22^, from which we extracted 20,310 CAD patients and 25,000 controls (Supplementary Table 3). From STARNET-based models, we identified 66 gene-tissue association pairs reaching Bonferroni-corrected significance (P<3.85e-6) in the ten CARDIoGRAMplusC4D cohorts. Of these, 19 also reached Bonferroni-corrected significance in the UKB data, which was significantly more than expected by chance (binomial test P = 0.00075), and 50 of 66 gene-tissue association pairs had directionally consistent effects (binomial test P =3.33e-5). We also found strong correlation of the effect sizes (ρ = 0.74; P = 1.3e-12; Supplementary Fig. 1A) indicating good overall reproducibility of the STARNET-based models.

From the GTEx-based models, 47 gene-tissue pairs reached Bonferroni-corrected significance (P<3.85e-6) in the ten CARDIoGRAMplusC4D cohorts, whereof 14 were significant also in UKB (binominal test P = 0.0079). Like the STARNET-based models, 39 of 44 significant gene-tissue association pairs had consistent direction of effects with a Pearsońs coefficient of 0.75 (P = 1.2e-9; Supplementary Fig. 1B). The slightly lower numbers of significant gene-tissue association pairs found in the GTEx models may be explained in that predicting models were based on: i) smaller numbers of genotype-expression pairs, ii) unlike STARNET, GTEx consist of apparently healthy tissues and iii) STARNET is a specific collection of CAD patients.

Next, we tested consistency of TWAS results between two reference-based models by comparing the results of a meta-analysis on all 11 genotyping data sets. We observed an average of 62% overlapping genes (Supplementary Table 1) and significant correlations of effect sizes (average Pearson’s coefficient ρ = 0.72; P < 1e-10; Supplementary Fig. 2). In the STARNET-based models, we identified 82 genes representing 129 gene-tissue pairs across seven tissues (P<3.85e-6). In the GTEx models, we identified 66 genes representing 106 gene-tissue pairs across eight tissues (P<3.85e-6). A total of 42 gene-tissue pairs were significant in both the STARNET- and GTEx-based models (Supplementary Fig. 3A). The overlapping genes were linearly consistent in both effect size (Pearson’s coefficient ρ = 0.99; P<2.2e-16) and -log_10_P (Pearsońs coefficient ρ = 0.82; P<4e-11) (Supplementary Fig. 3B). Overall, these results suggest, on the one hand, reasonable consistence between the two independent panels and, on the other hand, evidence for capturing complementary expression quantitative signals.

## Supplementary Tables

Supplementary Table 1. Statistics of nine tissues’ predictive models.

Supplementary Table 2. SNP priors of COR and TIB tissues.

Supplementary Table 3. 11 Genotype cohorts.

Supplementary Table 4. 114 TWAS genes list.

Supplementary Table 5. 53 TWAS genes have strong evidence of colocalized signals between GWAS and eQTL (PP4 > 0.55).

Supplementary Table 6. 96 known and 18 novel genes annotated by GWAS risk loci of CAD.

Supplementary Table 7. TWAS genes are enriched to CAD or related risk traits based on DisGeNET.

Supplementary Table 8. Pathways enriched by TWAS genes.

Supplementary Table 9. Association of TWAS genes’ damaging mutation with CAD and its binary risk traits.

Supplementary Table 10. Association of TWAS genes’ damaging mutation with quantitative risk traits of CAD.

Supplementary Table 11. Lead variants resided in the regions of novel genes were associated with lipid traits in human genotype data.

Supplementary Table 12. Expression-trait association statistics in mouse atherosclerosis model from HMDP.

Supplementary Table 13. Oligo sequences for gene editing.

## Supplementary Figures

**Supplementary Fig. 1.**
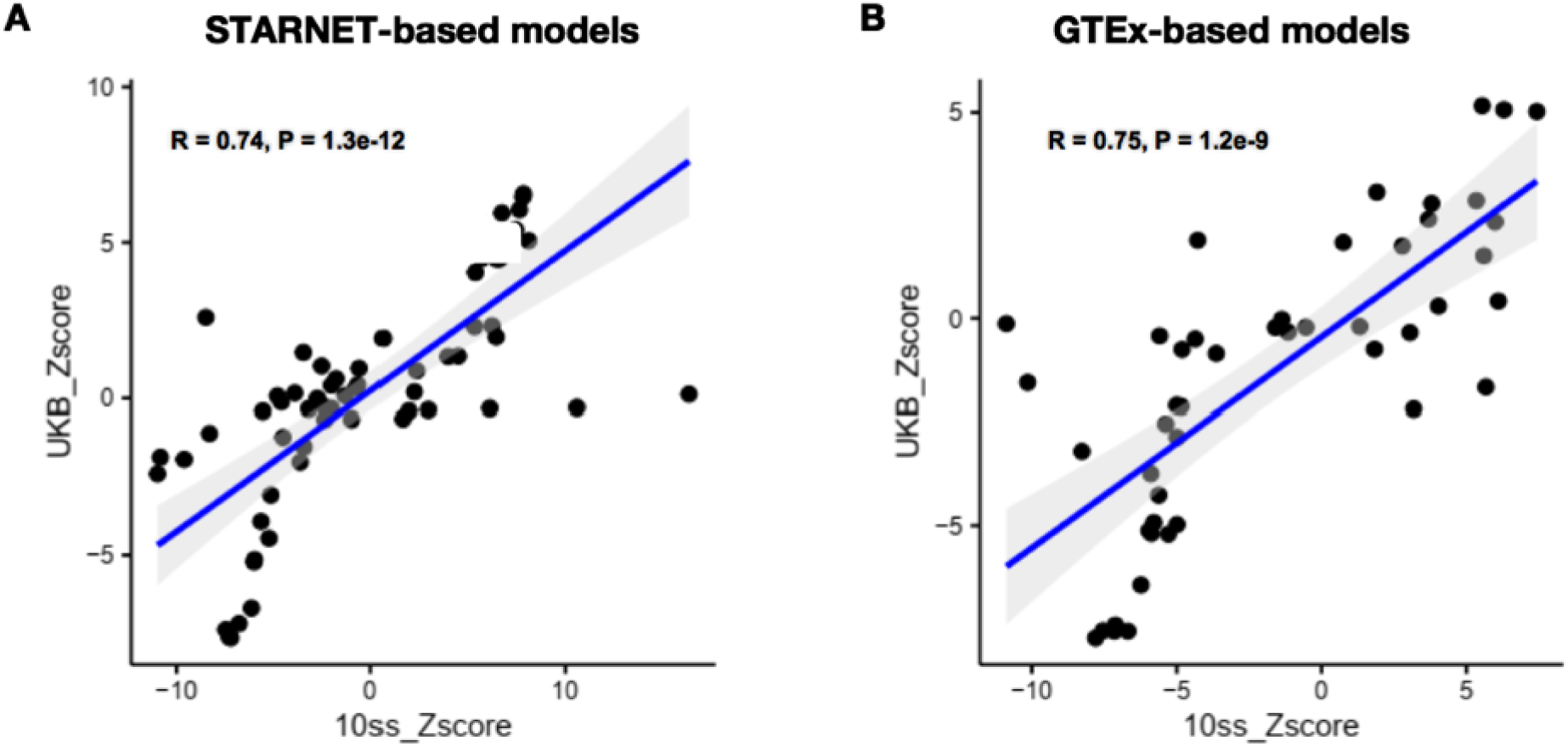
Reproducibility of TWAS results within two reference models. A) Reproducibility of STARNET-based models. B) Reproducibility of GTEx-based models. Ten CARDIoGRAMplusC4D cohorts (10ss) were used as the testing set, genotypes from UK Biobank (UKB) were the validating set.

**Supplementary Fig. 2.**
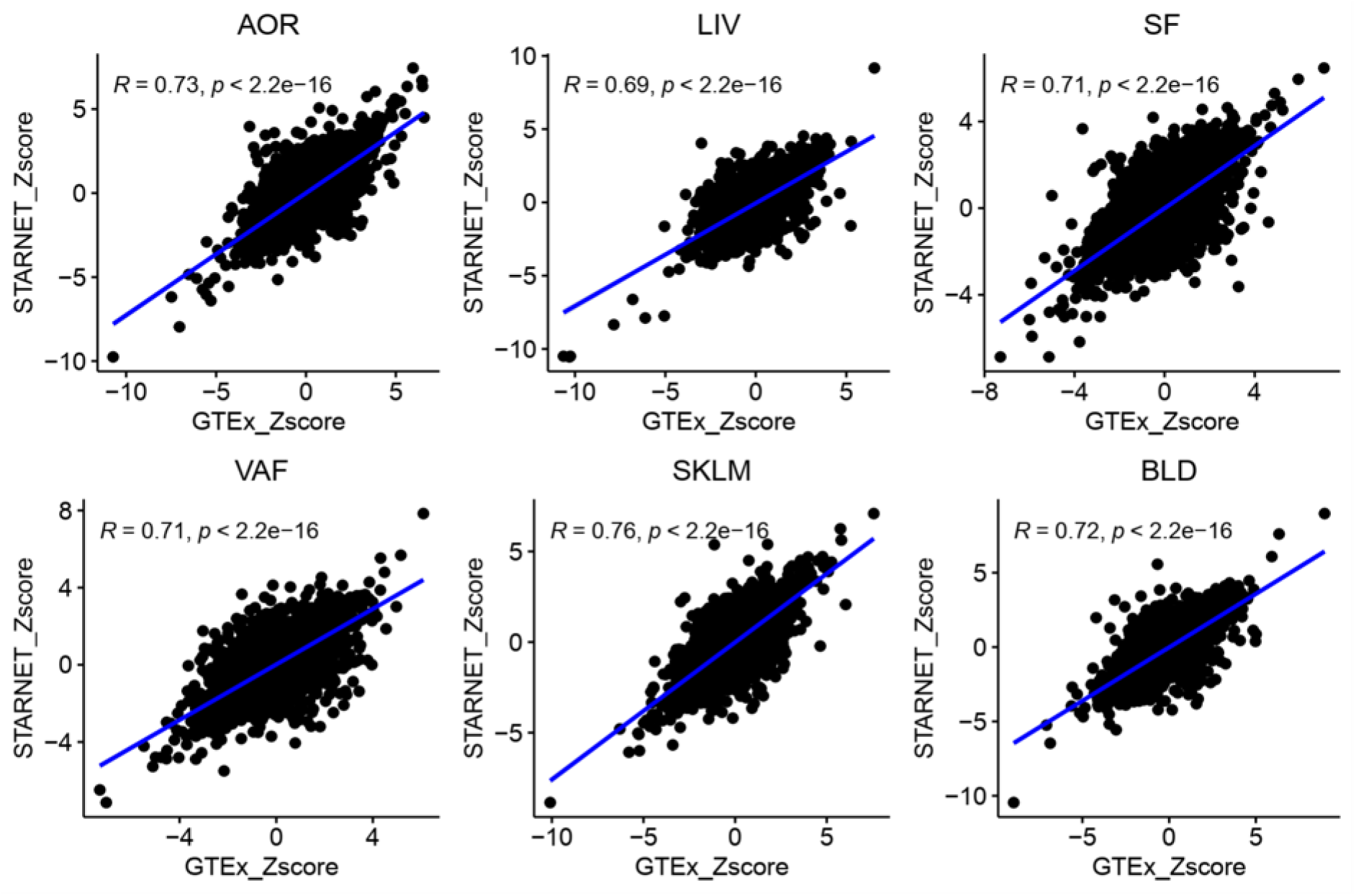
Associations of predicted expressions with CAD are consistent across tissues between STARNET- and GTEx-based models.

**Supplementary Fig. 3.**
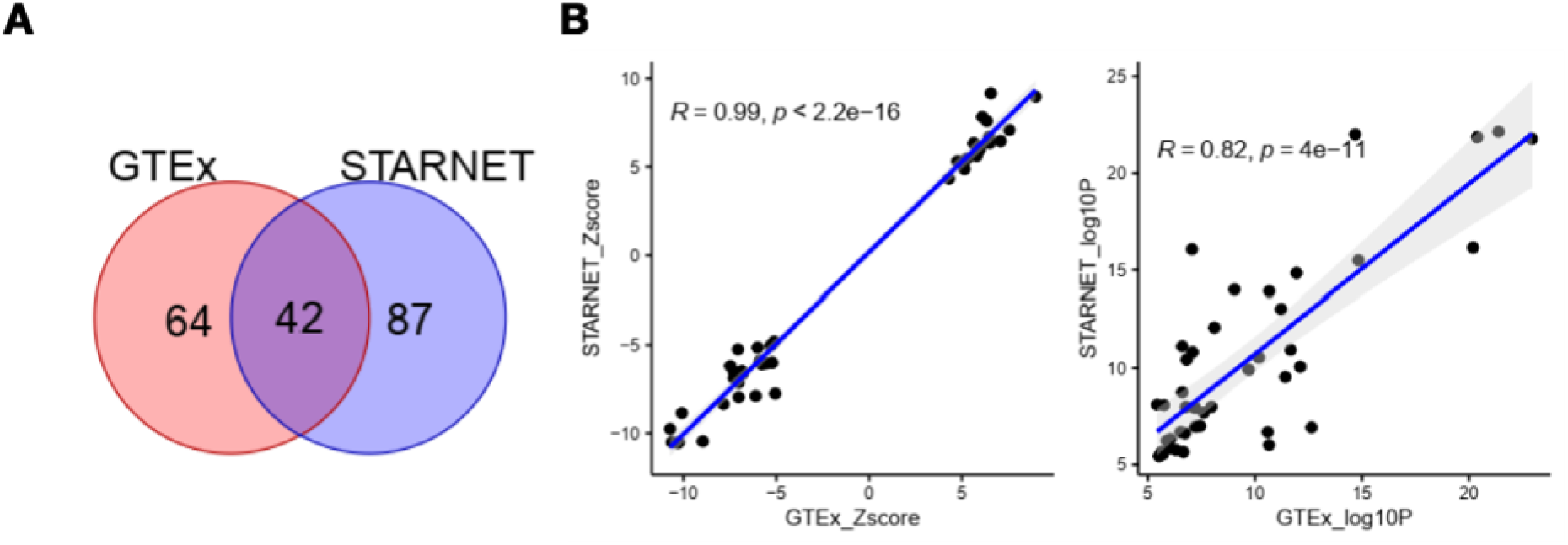
Comparation of TWAS results between two reference models. A) Venn diagram of transcriptome-wide significant gene-tissue pairs based on the two reference models. There are 42 overlapping gene-tissue pairs (34 genes). B) The effect sizes (left) and P values (right) of overlapping genes were consistent between the two reference-based models.

**Supplementary Fig. 4.**
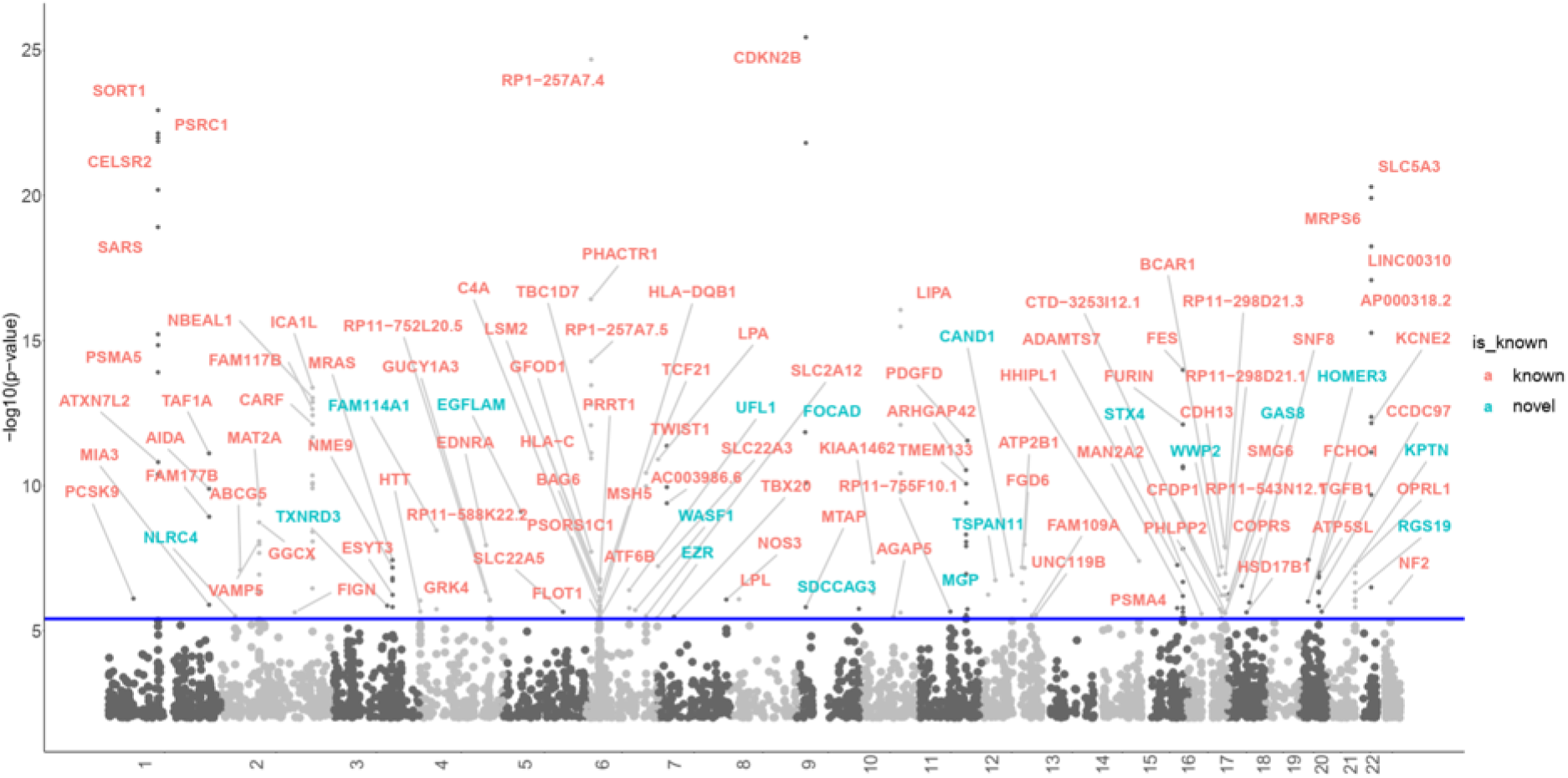
Manhattan plot of the transcriptome wide association study (TWAS). 114 TWAS genes are highlighted. The blue line marks P =3.85×10-6. Each point corresponds to an association test between a gene-tissue pair. TWAS genes residing in known GWAS loci were defined as known (red dots), otherwise defined as novel (blue dots).

**Supplementary Fig. 5.**
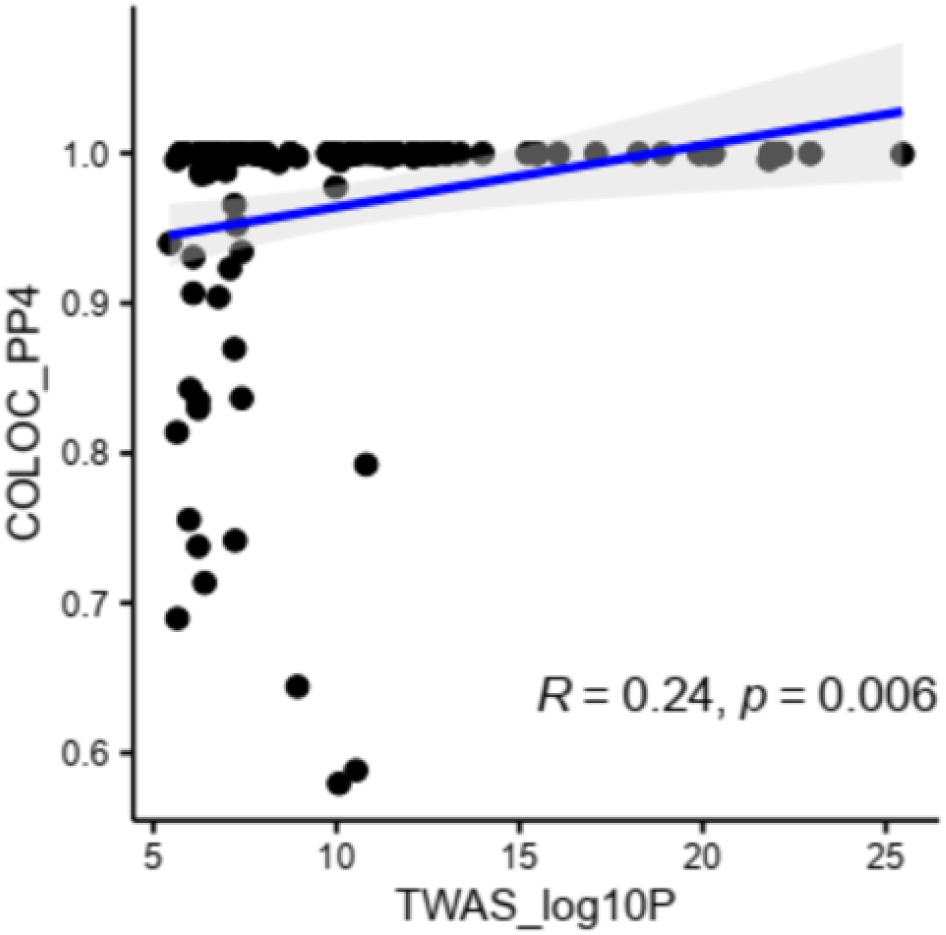
Positive correlation between TWAS and colocalization statistics. The log10P statistics of TWAS genes were positively correlated with PP4 (the posterior probabilities) statistics of colocalization analysis. Most TWAS genes have shared casual variants between GWAS and eQTL signals as their PP4 approaches 1.

**Supplementary Fig. 6.**
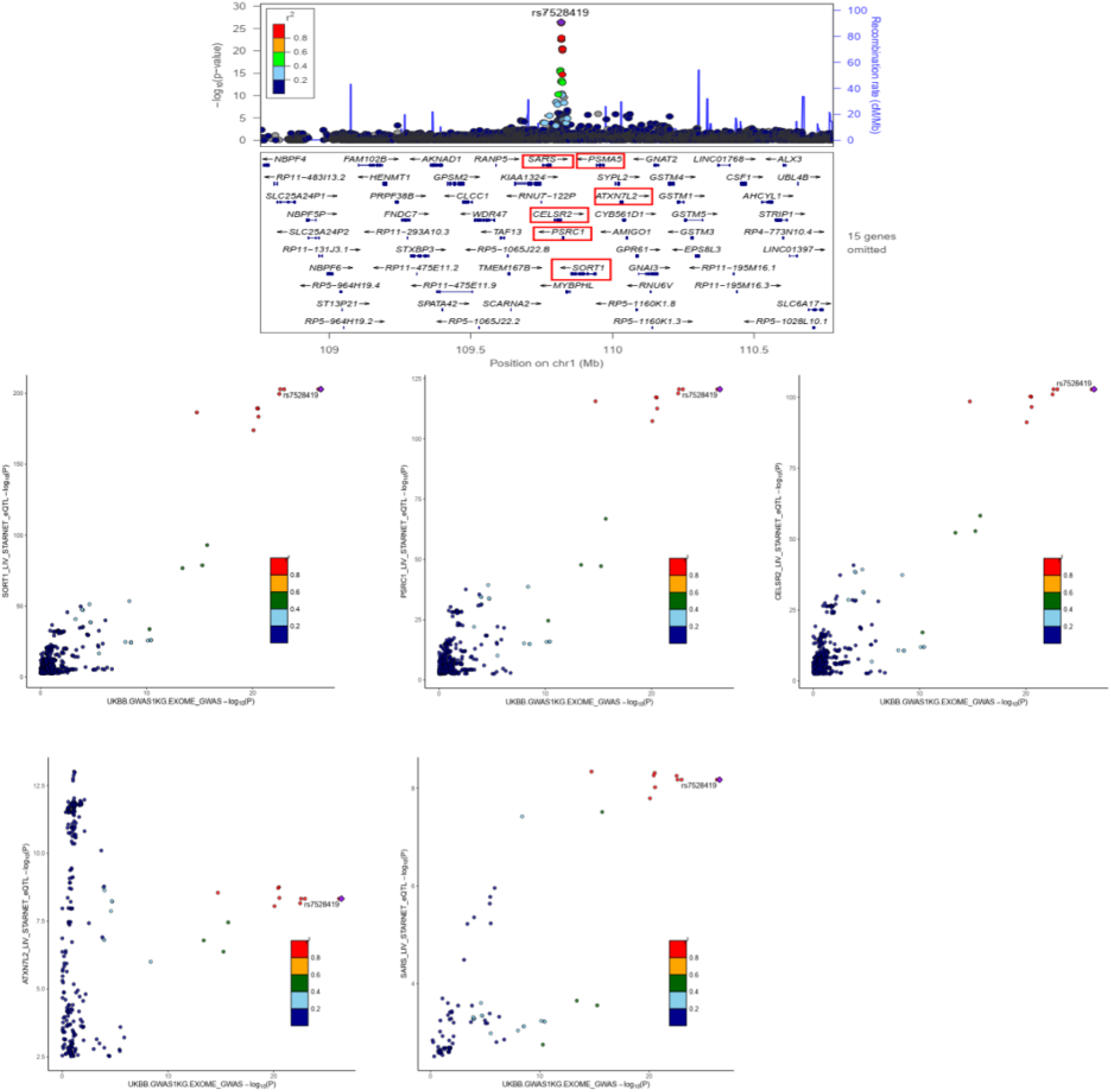
Colocalization signals in liver tissue at 1p13.3.

**Supplementary Fig. 7.**
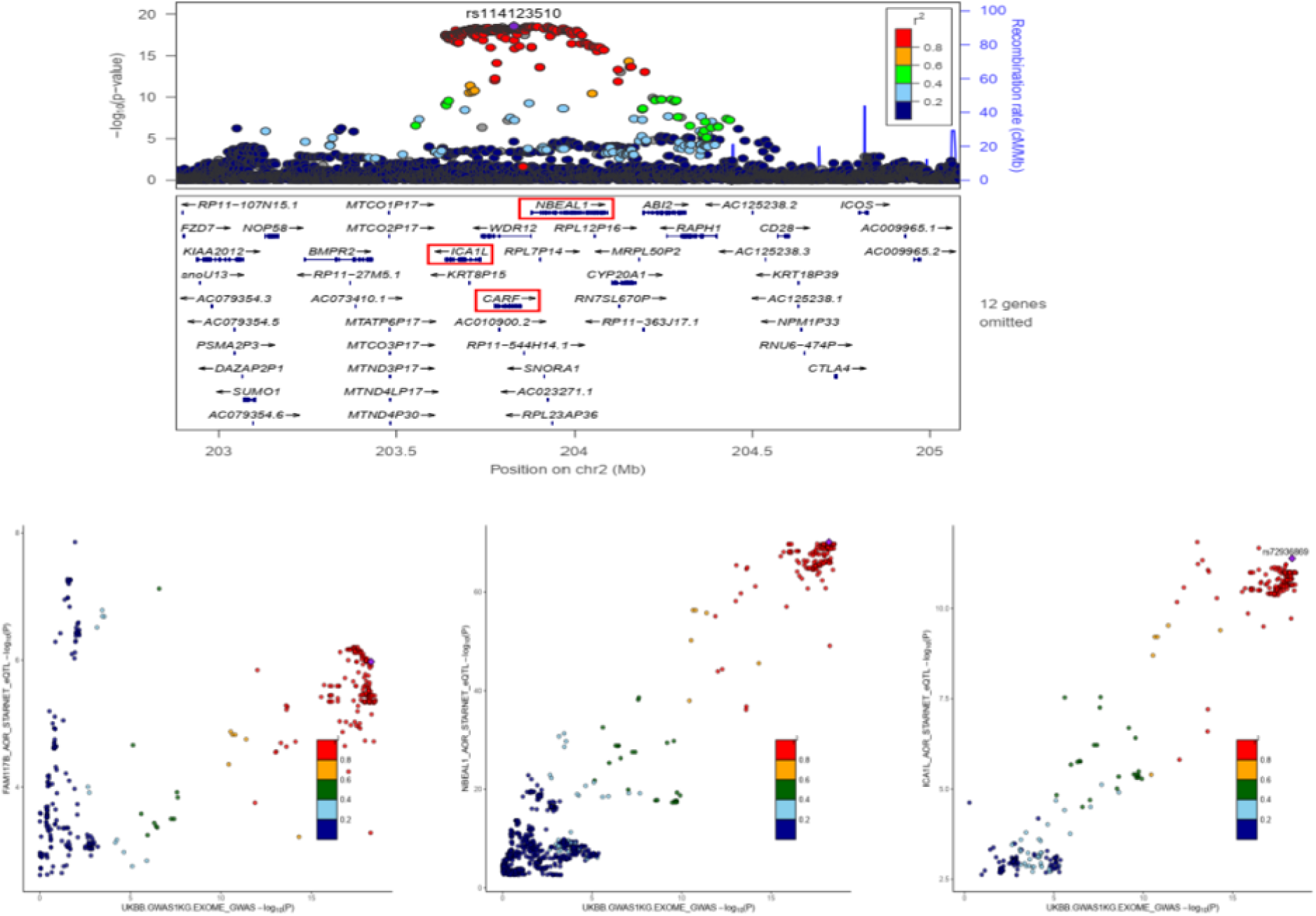
Colocalization signals in aorta tissue at 2p33.2.

**Supplementary Fig. 8.**
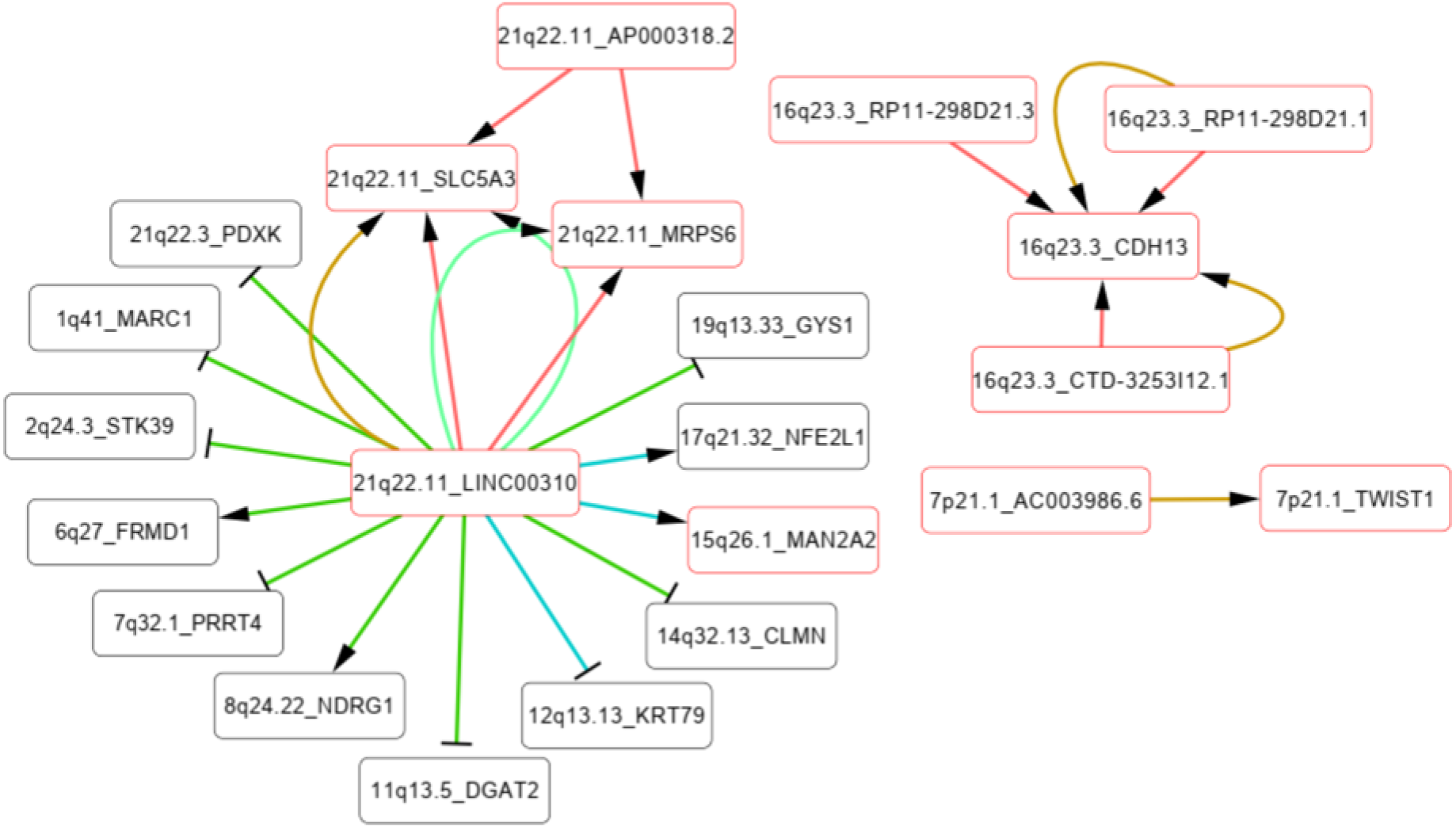
Co-expression network related to lncRNA genes. Coding genes with co-expression relationship with TWAS lncRNA genes are linked by arrow or T-line. Arrow suggests positive co-expression, and T-line suggests negative. TWAS genes are indicated in red frame. Tissues of gene co-expression are showed in difference edge colors as indicated.

**Supplementary Fig. 9.**
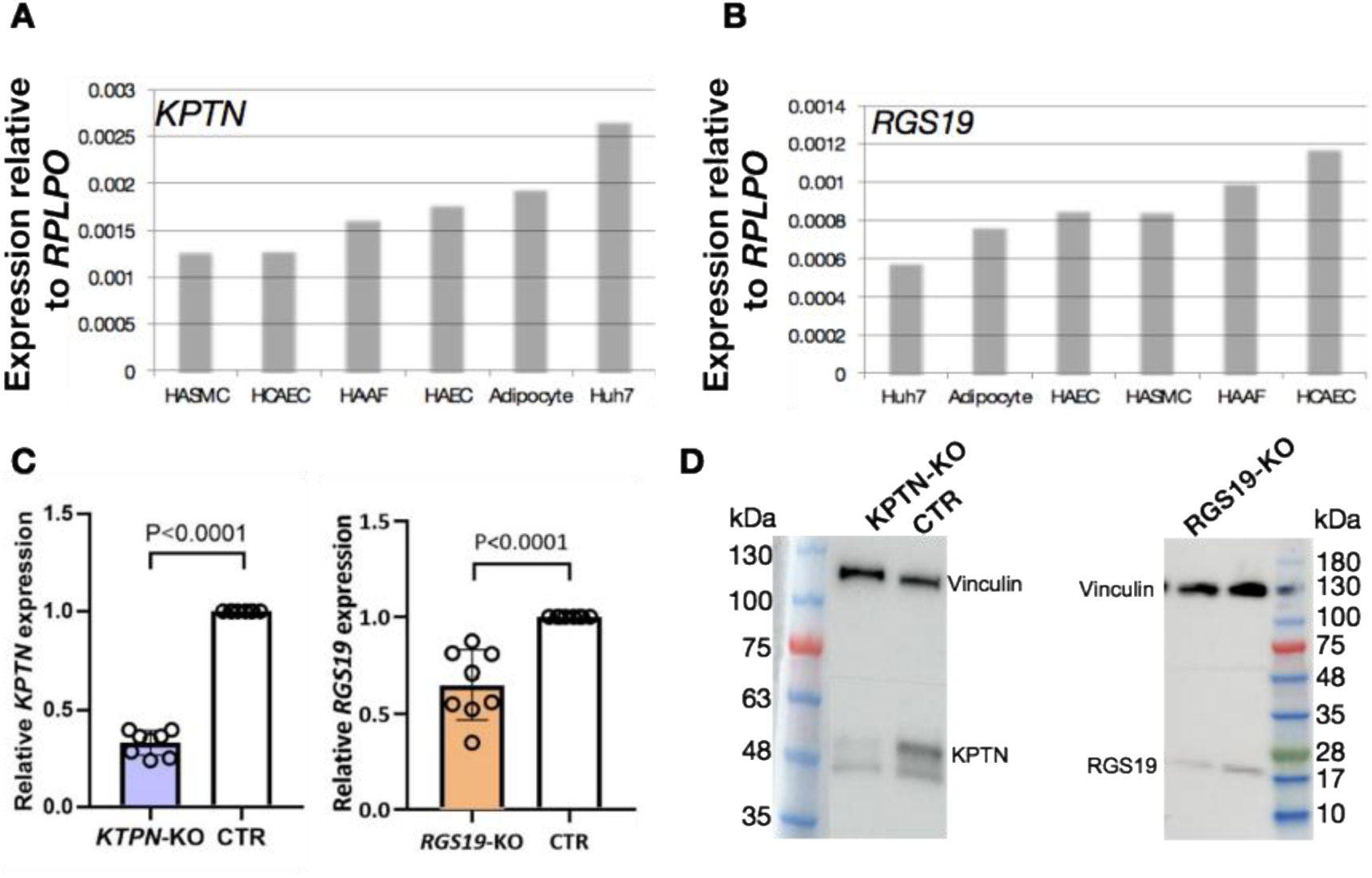
*KPTN* (A) and *RGS19* (B) expressions in multiple primary cells and cell lines. HASMC, human aorta smooth muscle cell; HCAEC, human coronary artery endothelium cell; HAAF, human aorta artery fibroblast; HAEC, human aorta endothelium cell and huh7, a human hepatoma cell line. (C) RNA levels of *KPTN* and *RGS19* were dramatically reduced in corresponding knockout lines (KO) in comparison to the control cell line (CTR), n=7. (D) The Western Blot image displays *KPTN* and *RGS19* reduction at protein level. Vinculin, 116kDa; *KPTN*, 48kDa; *RGS19*, 25kDa.

## Main References

1. Malakar, A. K. et al. A review on coronary artery disease, its risk factors, and therapeutics. J. Cell. Physiol. 234, 16812–16823 (2019).

2. Erdmann, J., Kessler, T., Munoz Venegas, L. & Schunkert, H. A decade of genome-wide association studies for coronary artery disease: The challenges ahead. Cardiovascular Research vol. 114 1241–1257 (2018).

3. Koyama, S. et al. Population-specific and trans-ancestry genome-wide analyses identify distinct and shared genetic risk loci for coronary artery disease. Nat. Genet. 52, 1169–1177 (2020).

4. Foroughi Asl, H., et al. Expression Quantitative Trait Loci Acting Across Multiple Tissues Are Enriched in Inherited Risk for Coronary Artery Disease. Circ. Cardiovasc. Genet. 8, 305–315 (2015).

5. Wild, P. S. et al. A Genome-Wide Association Study Identifies *LIPA* as a Susceptibility Gene for Coronary Artery Disease. Circ. Cardiovasc. Genet. 4, 403–412 (2011).

6. Vilne, B. & Schunkert, H. Integrating Genes Affecting Coronary Artery Disease in Functional Networks by Multi-OMICs Approach. Frontiers in Cardiovascular Medicine vol. 5 89 (2018).

7. Gamazon, E. R. et al. A gene-based association method for mapping traits using reference transcriptome data. Nat. Genet. 47, 1091–1098 (2015).

8. Wainberg, M. et al. Opportunities and challenges for transcriptome-wide association studies. Nat. Genet. 51, 592–599 (2019).

9. Zhang, W. et al. Integrative transcriptome imputation reveals tissue-specific and shared biological mechanisms mediating susceptibility to complex traits. Nat. Commun. 10, 1–13 (2019).

10. Franzén, O. et al. Cardiometabolic risk loci share downstream cis- and trans-gene regulation across tissues and diseases. Science (80-. ). 353, 827–830 (2016).

11. Lonsdale, J. et al. The Genotype-Tissue Expression (GTEx) project. Nature Genetics vol. 45 580–585 (2013).

12. Samani, N. J. et al. Genomewide Association Analysis of Coronary Artery Disease. N. Engl. J. Med. 357, 443–453 (2007).

13. Erdmann, J. et al. New susceptibility locus for coronary artery disease on chromosome 3q22.3. Nat. Genet. 41, 280–282 (2009).

14. Erdmann, J. et al. Genome-wide association study identifies a new locus for coronary artery disease on chromosome 10p11.23. Eur. Heart J. 32, 158–168 (2011).

15. Nikpay, M. et al. A comprehensive 1000 Genomes-based genome-wide association meta-analysis of coronary artery disease. Nat. Genet. 47, 1121–1130 (2015).

16. Stitziel, N. O. et al. Inactivating mutations in NPC1L1 and protection from coronary heart disease. N. Engl. J. Med. 371, 2072–2082 (2014).

17. Nelson, C. P. et al. Association analyses based on false discovery rate implicate new loci for coronary artery disease. Nat. Genet. 49, 1385–1391 (2017).

18. Li, L., Pang, S., Zeng, L., Güldener, U. & Schunkert, H. Genetically determined intelligence and coronary artery disease risk. Clin. Res. Cardiol. (2020) doi:10.1007/s00392-020-01721-x.

19. Burton, P. R. et al. Genome-wide association study of 14,000 cases of seven common diseases and 3,000 shared controls. Nature 447, 661–678 (2007).

20. Winkelmann, B. R. et al. Rationale and design of the LURIC study - A resource for functional genomics, pharmacogenomics and long-term prognosis of cardiovascular disease. Pharmacogenomics 2, (2001).

21. Anderson, C. D. et al. Genome-wide association of early-onset myocardial infarction with single nucleotide polymorphisms and copy number variants. Nat. Genet. 478, 103–109 (2015).

22. Bycroft, C. et al. The UK Biobank resource with deep phenotyping and genomic data. Nature 562, 203–209 (2018).

23. Giambartolomei, C. et al. Bayesian Test for Colocalisation between Pairs of Genetic Association Studies Using Summary Statistics. PLoS Genet. 10, (2014).

24. Piñero, J. et al. The DisGeNET knowledge platform for disease genomics: 2019 update. Nucleic Acids Res. 48, D845–D855 (2020).

25. Harris, M. A. et al. The Gene Oncology (GO) database and informatics resource. Nucleic Acids Res. 32, D258–D261 (2004).

26. Kanehisa, M. & Goto, S. KEGG: Kyoto Encyclopedia of Genes and Genomes. Nucleic Acids Research vol. 28 27–30 (2000).

27. Croft, D. et al. Reactome: A database of reactions, pathways and biological processes. Nucleic Acids Res. 39, D691–D697 (2011).

28. Slenter, D. N. et al. WikiPathways: A multifaceted pathway database bridging metabolomics to other omics research. Nucleic Acids Res. 46, D661–D667 (2018).

29. Bjørklund, G. et al. The Role of Matrix Gla Protein (MGP) in Vascular Calcification. Curr. Med. Chem. 27, 1647–1660 (2019).

30. Okla, M. et al. Ellagic acid modulates lipid accumulation in primary human adipocytes and human hepatoma Huh7 cells via discrete mechanisms. J. Nutr. Biochem. 26, 82–90 (2015).

31. Li, C. et al. Matrix Gla protein regulates adipogenesis and is serum marker of visceral adiposity. Adipocyte 9, 68–76 (2020).

32. Borborema, M. E. de A., Crovella, S., Oliveira, D. & de Azevêdo Silva, J. Inflammasome activation by NLRP1 and NLRC4 in patients with coronary stenosis. Immunobiology 225, 151940 (2020).

33. Alehashemi, S. & Goldbach-Mansky, R. Human Autoinflammatory Diseases Mediated by NLRP3-, Pyrin-, NLRP1-, and NLRC4-Inflammasome Dysregulation Updates on Diagnosis, Treatment, and the Respective Roles of IL-1 and IL-18. Frontiers in Immunology vol. 11 1840 (2020).

34. Lusis, A. J. et al. The hybrid mouse diversity panel: A resource for systems genetics analyses of metabolic and cardiovascular traits. Journal of Lipid Research vol. 57 925–942 (2016).

35. Tiwari, S. & Siddiqi, S. A. Intracellular trafficking and secretion of VLDL. Arteriosclerosis, Thrombosis, and Vascular Biology vol. 32 1079–1086 (2012).

36. Flannick, J. et al. Exome sequencing of 20,791 cases of type 2 diabetes and 24,440 controls. Nature 570, 71–76 (2019).

37. Wolfson, R. L. et al. KICSTOR recruits GATOR1 to the lysosome and is necessary for nutrients to regulate mTORC1. Nature 543, 438–442 (2017).

38. Oishi, Y. et al. SREBP1 Contributes to Resolution of Pro-inflammatory TLR4 Signaling by Reprogramming Fatty Acid Metabolism. Cell Metab. 25, 412–427 (2017).

39. Tso, P. H., Yung, L. Y., Wang, Y. & Wong, Y. H. RGS19 stimulates cell proliferation by deregulating cell cycle control and enhancing Akt signaling. Cancer Lett. 309, 199–208 (2011).

40. Sangphech, N., Osborne, B. A. & Palaga, T. Notch signaling regulates the phosphorylation of Akt and survival of lipopolysaccharide-activated macrophages via regulator of G protein signaling 19 (RGS19). Immunobiology 219, 653–660 (2014).

41. Jansen, H., Samani, N. J. & Schunkert, H. Mendelian randomization studies in coronary artery disease. European Heart Journal vol. 35 1917–1924 (2014).

42. McGough, I. J. et al. Identification of molecular heterogeneity in SNX27-retromermediated endosome-to-plasma-membrane recycling. J. Cell Sci. 127, 4940–4953 (2014).

43. Sixt, S. et al. Long-but not short-term multifactorial intervention with focus on exercise training improves coronary endothelial dysfunction in diabetes mellitus type 2 and coronary artery disease. Eur. Heart J. 31, 112–119 (2010).

44. Arvind, P., Nair, J., Jambunathan, S., Kakkar, V. V. & Shanker, J. CELSR2-PSRC1-SORT1 gene expression and association with coronary artery disease and plasma lipid levels in an Asian Indian cohort. J. Cardiol. 64, 339–346 (2014).

45. Coding Variation in ANGPTL4, LPL, and SVEP1 and the Risk of Coronary Disease. N. Engl. J. Med. 374, 1134–1144 (2016).

46. Tsutsumi, K. Lipoprotein Lipase and Atherosclerosis. Curr. Vasc. Pharmacol. 1, 11–17 (2003).

## Methods References

1. Zhang, W. et al. Integrative transcriptome imputation reveals tissue-specific and shared biological mechanisms mediating susceptibility to complex traits. Nat. Commun. 10, 1–13 (2019).

2. Franzén, O. et al. Cardiometabolic risk loci share downstream cis- and trans-gene regulation across tissues and diseases. Science (80-. ). 353, 827–830 (2016).

3. Aguet, F. et al. Genetic effects on gene expression across human tissues. Nature 550, 204–213 (2017).

4. Li, Y. I. et al. RNA splicing is a primary link between genetic variation and disease. Science (80-. ). 352, 600–604 (2016).

5. Bernstein, B. E. et al. The NIH roadmap epigenomics mapping consortium. Nature Biotechnology vol. 28 1045–1048 (2010).

6. Samani, N. J. et al. Genomewide association analysis of coronary artery disease. N. Engl. J. Med. 357, 443–453 (2007).

7. Erdmann, J. et al. New susceptibility locus for coronary artery disease on chromosome 3q22.3. Nat. Genet. 41, 280–282 (2009).

8. Erdmann, J. et al. Genome-wide association study identifies a new locus for coronary artery disease on chromosome 10p11.23. Eur. Heart J. 32, 158–168 (2011).

9. Nikpay, M. et al. A comprehensive 1000 Genomes-based genome-wide association meta-analysis of coronary artery disease. Nat. Genet. 47, 1121–1130 (2015).

10. Stitziel, N. O. et al. Inactivating mutations in NPC1L1 and protection from coronary heart disease. N. Engl. J. Med. 371, 2072–2082 (2014).

11. Nelson, C. P. et al. Association analyses based on false discovery rate implicate new loci for coronary artery disease. Nat. Genet. 49, 1385–1391 (2017).

12. Li, L., Pang, S., Zeng, L., Güldener, U. & Schunkert, H. Genetically determined intelligence and coronary artery disease risk. Clin. Res. Cardiol. 1–9 (2020) doi:10.1007/s00392-020-01721-x.

13. Burton, P. R. et al. Genome-wide association study of 14,000 cases of seven common diseases and 3,000 shared controls. Nature 447, 661–678 (2007).

14. Winkelmann, B. R. et al. Rationale and design of the LURIC study - A resource for functional genomics, pharmacogenomics and long-term prognosis of cardiovascular disease. Pharmacogenomics 2, (2001).

15. Anderson, C. D. et al. Genome-wide association of early-onset myocardial infarction with single nucleotide polymorphisms and copy number variants. Nat. Genet. 478, 103–109 (2015).

16. Bycroft, C. et al. The UK Biobank resource with deep phenotyping and genomic data. Nature 562, 203–209 (2018).

17. Kohl, M., Wiese, S. & Warscheid, B. Cytoscape: software for visualization and analysis of biological networks. Methods Mol. Biol. 696, 291–303 (2011).

18. Giambartolomei, C. et al. Bayesian Test for Colocalisation between Pairs of Genetic Association Studies Using Summary Statistics. PLoS Genet. 10, (2014).

19. Erdmann, J., Kessler, T., Munoz Venegas, L. & Schunkert, H. A decade of genome-wide association studies for coronary artery disease: The challenges ahead. Cardiovascular Research vol. 114 1241–1257 (2018).

20. Koyama, S. et al. Population-specific and trans-ancestry genome-wide analyses identify distinct and shared genetic risk loci for coronary artery disease. Nat. Genet. 52, 1169–1177 (2020).

21. De Leeuw, C. A., Mooij, J. M., Heskes, T. & Posthuma, D. MAGMA: Generalized Gene-Set Analysis of GWAS Data. PLoS Comput Biol 11, 1004219 (2015).

22. Bindea, G. et al. ClueGO: A Cytoscape plug-in to decipher functionally grouped gene ontology and pathway annotation networks. Bioinformatics 25, 1091–1093 (2009).

23. Harris, M. A. et al. The Gene Oncology (GO) database and informatics resource. Nucleic Acids Res. 32, D258–D261 (2004).

24. Kanehisa, M. & Goto, S. KEGG: Kyoto Encyclopedia of Genes and Genomes. Nucleic Acids Research vol. 28 27–30 (2000).

25. Croft, D. et al. Reactome: A database of reactions, pathways and biological processes. Nucleic Acids Res. 39, D691–D697 (2011).

26. Slenter, D. N. et al. WikiPathways: A multifaceted pathway database bridging metabolomics to other omics research. Nucleic Acids Res. 46, D661–D667 (2018).

27. Piñero, J. et al. The DisGeNET knowledge platform for disease genomics: 2019 update. Nucleic Acids Res. 48, D845–D855 (2020).

28. Van Hout, C. V. et al. Exome sequencing and characterization of 49,960 individuals in the UK Biobank. Nature 586, 749–756 (2020).

29. Landrum, M. J. et al. ClinVar: improvements to accessing data. Nucleic Acids Res. 48, 835–844 (2019).

30. McLaren, W. et al. The Ensembl Variant Effect Predictor. Genome Biol. 17, (2016).

31. Karczewski, K. J. et al. The mutational constraint spectrum quantified from variation in 141,456 humans. Nature 581, 434–443 (2020).

32. Dong, C. et al. Comparison and integration of deleteriousness prediction methods for nonsynonymous SNVs in whole exome sequencing studies. Hum. Mol. Genet. 24, 2125–2137 (2015).

33. Chang, C. C. et al. Second-generation PLINK: rising to the challenge of larger and richer datasets. Gigascience 4, 7 (2015).

35. Bennett, B. J. et al. Genetic Architecture of Atherosclerosis in Mice: A Systems Genetics Analysis of Common Inbred Strains. PLoS Genet. 11, 1005711 (2015).

